# Evolutionary dynamics of the Timopheevii wheat lineage

**DOI:** 10.64898/2026.07.20.739399

**Authors:** Laxman Adhikari, Emile Cavalet-Giorsa, Michael Abrouk, Dal-Hoe Koo, Samara Correia de Lemos, Thomas Lux, Georg Haberer, Vitaly Portnoy, Adil Salhi, Xin Gao, Péter Mikó, István Molnár, Nathalie Rodde, W. John Raupp, Hakan Özkan, Manuel Spannagl, Nadia Kamal, Assaf Distelfeld, Jesse Poland, Simon G. Krattinger

## Abstract

Wheat, the most widely cultivated crop globally, includes multiple species with varying ploidy levels^1^. Globally cultivated species such as pasta and bread wheat belong to the Emmer lineage. In contrast, the Timopheevii lineage is of negligible agricultural significance^2^, yet it harbours genetic diversity of potential value for wheat improvement. Here, we present genomic resources for the Timopheevii wheat lineage. Our analyses uncovered extensive homoeologous exchange and genome instability in zhukovsky’s wheat, the hexaploid representative of the Timopheevii lineage. Population genomic analyses reveal a mosaic-like haplotype composition of domesticated tetraploid timopheev’s wheat, suggesting it was once cultivated across a broader geographic range. This is further supported by the discovery of ‘fossil’ haplotype segments in bread wheat landraces, likely derived from now-extinct domesticated timopheev’s wheat populations. By examining the Timopheevii lineage, we provide new insights into the evolutionary dynamics of wheat, discovering previously underexplored and potentially useful genetic diversity for wheat improvement.

## Main

Wheat is one of the most successful and most widely cultivated crops globally, accounting for around 20% of both protein and caloric intake. Current yield gains, however, are projected to be insufficient to meet future wheat demands^3,4^. Capturing, mapping, and understanding the genetic diversity within the gene pools of wheat and its wild relatives has been identified as a critical component to breed high-yielding and climate-resilient wheat varieties^3^.

There are two main wheat lineages with independent evolutionary origins. Both lineages comprise multiple wild and domesticated species with different ploidy levels^1^. The Emmer lineage has the basic genome composition BBAA and arose through a hybridization between *Triticum urartu* and a relative of *Aegilops speltoides*. The Timopheevii lineage has the basic genome composition GGA^t^A^t^, with the G genome likely originating from a different *Ae. speltoides* population^5^. Presently, the only economically relevant wheats on a global scale are domesticated species within the Emmer lineage, namely tetraploid durum (*T. turgidum* ssp. *durum*) and hexaploid bread wheat (*T. aestivum* ssp. *aestivum*). These two wheat species account for virtually all of the global wheat production. The domesticated species within the Timopheevii lineage, tetraploid timopheev’s wheat (*T. timopheevii* ssp. *timopheevii;* 2*n =* 4*x=* 28; GGA^t^A^t^ genome) and hexaploid zhukovsky’s wheat (*T. zhukovskyi;* 2*n =* 6*x =* 42, GGA^t^A^t^A^m^A^m^ genome)^6^, on the other hand, do not play a major role in global wheat production. Timopheev’s wheat cultivation is limited to Transcaucasia, particularly Georgia, although its distribution range might have been broader^1,2,7^. The allohexaploid *T. zhukovskyi* has only been reported in a restricted geographical area in western Georgia and only in mixed fields containing diploid einkorn wheat (*T. monococcum;* 2*n =* 2*x =* 14; A^m^A^m^ genome) and *T. timopheevii*^1^. It has thus been suggested that *T. zhukovskyi* arose through a hybridization between einkorn and timopheev’s wheat. The difference in importance of the Timopheevii and Emmer lineage is intriguing: Why did the Emmer wheat lineage become globally dominant in modern times, while the closely related Timopheevii lineage did not? Has the Timopheevii lineage been more relevant for agriculture in the past?

Here, we generate and analyse comprehensive genomic resources for the Timopheevii wheat lineage, including chromosome-scale assemblies of wild and domesticated timopheev’s wheat, two contig-level assemblies of zhukovsky’s wheat, and whole-genome sequencing data of a timopheev’s wheat diversity panel. Our analyses provide insights into genome dynamics in the Timopheevii lineage, revealing its legacy on bread wheat improvement.

## Results

### Timopheevii wheat lineage assemblies

We produced chromosome-scale assemblies for two wild (*T. timopheevii* ssp. *armeniacum,* TA877 = PI 427325, northern Iraq; and KU-1984B, Türkiye) and one domesticated (*T. timopheevii* ssp. *timopheevii;* TA2804) tetraploid timopheev’s wheat accession (Fig. 1a). Northern Iraq has been suggested as the centre of diversity for wild timopheev’s wheat^2^. The three assemblies, constructed from PacBio HiFi reads, Hi-C data, and optical maps (for TA877 and TA2804), had final assembly sizes from 9.55 to 9.72 Gb, with contig N50s ranging from 33.0 to 42.7 Mb (Extended Data Table 1, Supplementary Fig. 1). *De novo* annotation^8^ predicted 87,967 and 88,243 gene models for TA877 and TA2804, respectively (Extended Data Table 1). For KU-1984B, 88,257 gene models were projected based on the other two *T. timopheevii de novo* annotations. Comparative analysis revealed a major reciprocal translocation between chromosomes 7A^t^ and 4G between TA877 and the other two accessions (Fig. 1a, b), with the translocation breakpoint located in the functional centromere region (Fig. 1b, Supplementary Fig. 2). Thus, the 7A^t^ :4G translocation in TA877 appears to be a whole-arm reciprocal translocation (Robertsonian-like translocation), while the 4G chromosome of TA2804 also shows a translocation involving chromosome 5A (Fig. 1a).

**Fig. 1.**
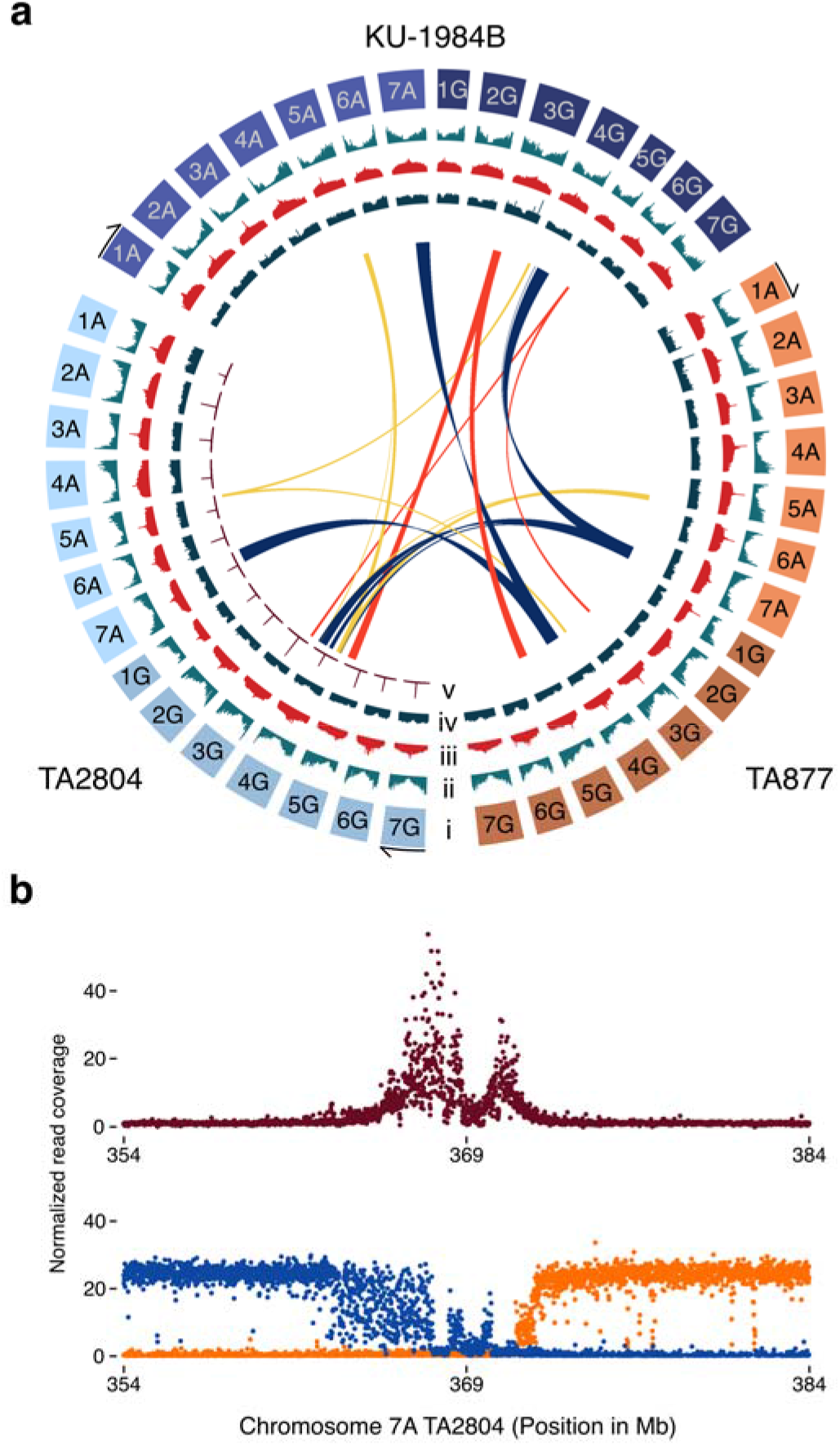
*T. timopheevii* genome structure and functional features. **a,** Circos plot showing synteny and the major translocation among the assemblies of the wild *T. timopheevii* accessions TA877 and KU-1984B, and the domesticated *T. timopheevii* accession TA2804. The tracks display the functional and structural features of the three assemblies. The number and length of chromosomes, with arrow indicating chromosome orientation (i), high-confidence gene density along the pseudomolecules (ii), density of Gypsy LTR retrotransposons (iii), density of Copia LTR retrotransposons (iv), and CENH3 ChIP-Seq read coverage for TA2804, with peaks indicating functional centromere positions (v). The inner circle highlights syntenic blocks involved in the major reciprocal translocations among the domesticated and two wild *T. timopheevii*. **b,** Read coverage plot along chromosome 7A of the domesticated *T. timopheevii* accession TA2804. The top panel shows CENH3-ChIP-Seq read coverage in dark red colour. In the lower panel, reads from the wild accession TA877 originating from chromosome 7A are shown in blue colour and reads originating from chromosome 4G are shown in orange colour. The translocation breakpoint lies within the functional centromere region.

In addition to the three chromosome-scale *T. timopheevii* assemblies, we produced contig-level assemblies for two *T. zhukovskyi* accessions, TA11273 (= PI 352552) and TA11274 (= PI 352553). For each accession we generated ∼260 Gb of PacBio HiFi data, corresponding to a ∼18-fold coverage. Surprisingly, the total assembly sizes of 12.0 and 11.2 Gb for TA11273 and TA11274 were considerably shorter than the expected ∼14.7 Gb for a hexaploid *T. zhukovskyi* assembly (5.2 Gb for einkorn + 9.5 Gb for *T. timopheevii,* Supplementary Table 1). Using identity-by-state analyses^9,10^, we could assign 4.19 Gb of the TA11273 assembly to einkorn wheat and 7.87 Gb to *T. timopheevii*. For TA11274, 3.22 Gb could be assigned to einkorn wheat and 8.04 Gb to *T. timopheevii* (Supplementary Table 1). The reduced assembly lengths could indicate homoeologous exchange between the A^m^ and A^t^ genomes in *T. zhukovskyi,* resulting in ‘collapsed’ genomic segments that have become fixed across the four A genome chromosomes (Fig. 2a, b, Extended Data Fig. 1a). Homoeologous exchange can result in the replacement of segments of one subgenome with segments from the other subgenome. Haplotypes of these segments can thus increase from two copies to three or four copies (or decrease to zero copies), resembling the chromosome composition of an autopolyploid species^11^. Mapping raw PacBio reads to the *T. zhukovskyi* contig-level assemblies revealed a bimodal coverage distribution (Extended Data Fig. 1b), with peaks at the expected ∼18-fold depth and a second peak at ∼36-fold. The second peak corresponds to regions identical across all four A-genome chromosomes that collapsed during assembly. Previous cytogenetic analyses revealed variation in *T. zhukovskyi* karyotypes, which could be caused by homoeologous exchange between the A^t^ and A^m^ genomes^12^.

**Fig. 2.**
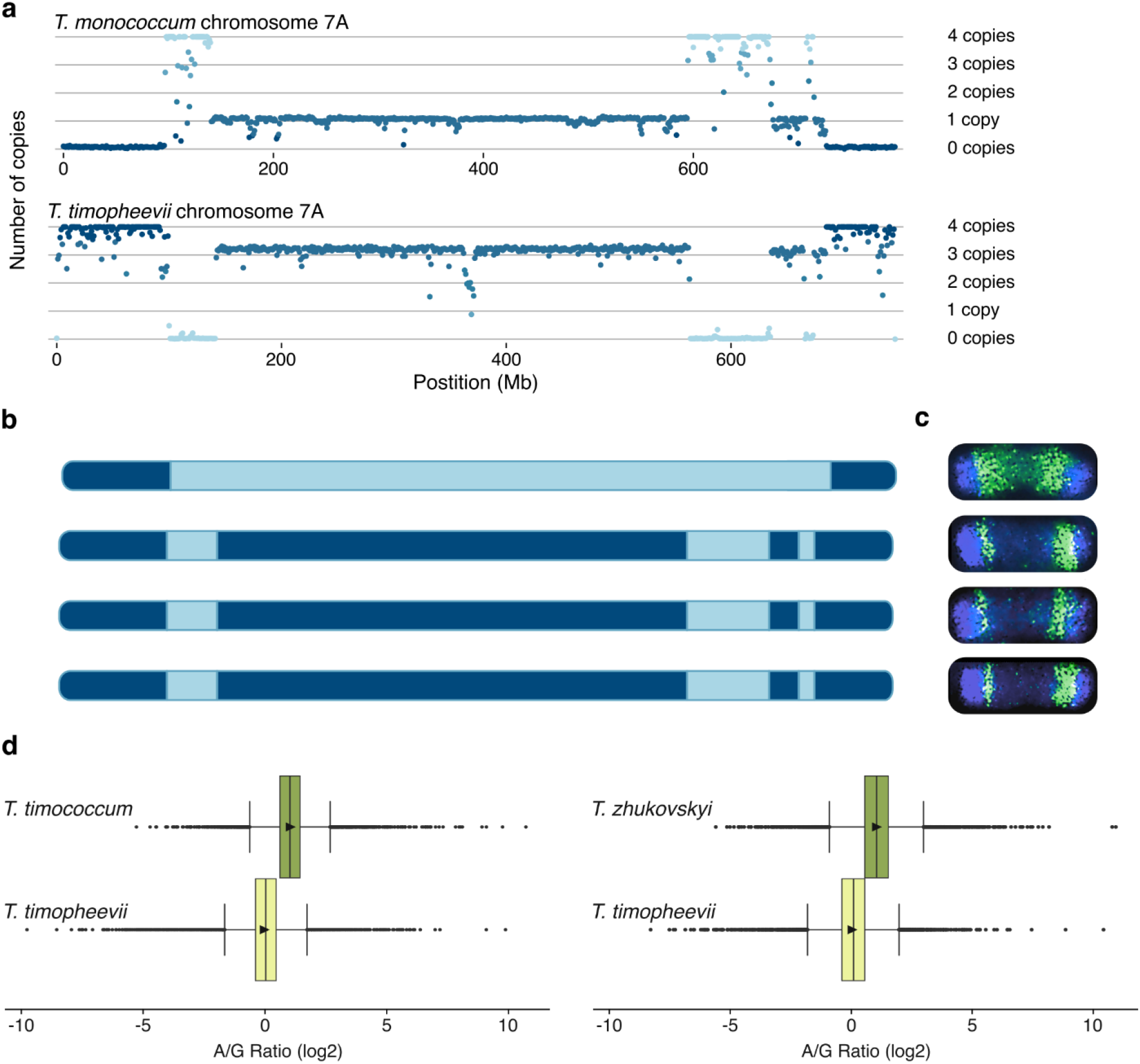
Homoeologous exchange in hexaploid zhukovsky’s wheat. **a,** Normalized read counts corresponding to chromosomes 7A^m^ and 7A^t^ in *T. zhukovskyi* plant TA11273_P4. **b,** Schematic representation of the four copies of chromosome 7A in *T. zhukovskyi* plant TA11273_P4. Segments in dark blue colour originate from *T. timopheevii* and segments shown in light blue colour from *T. monococcum*. **c,** Oligo painting on mitotic metaphase chromosomes of *T. zhukovskyi* plant TA11273_P4. Probes designed on chromosome 7A of einkorn wheat are shown in green colour and probes designed on chromosome 7A of *T. timopheevii* in blue colour. The blue oligo-FISH signals represent a pseudo-colour (original colour was red). **d,** A-to-G gene expression ratios across tetraploid and hexaploid representative of the Timopheevii wheat lineage. The results are based on 11,607 single-copy dyads (in the tetraploid) and triads (in the hexaploid). Boxes show the interquartile range (IQR) and the median of the A-to-G gene expression ratios for triads and dyads. Whiskers show the range of the data within 1.5 times the IQR and dots represent outliers that fall outside the whiskers, indicating extreme values in the data. The triangle represent the mean (average) value of the A-to-G ratio for each group.

### Genome instability in *T. zhukovskyi*

To further investigate the potential genome instability in *T. zhukovskyi*, we investigated 19 individual plants representing *T. zhukovskyi* accessions TA11273, TA11274 and TA2610. Whole-genome sequencing of these plants revealed that the *T*. *zhukovskyi* individuals are essentially redundant genotypes (>98.5% allele matching), supporting previous notions that all *T. zhukovskyi* accessions maintained in *ex situ* collections are derived from one original genotype^2^ (Extended Data Fig. 2). Similarly, the mean genome-wide nucleotide diversity (π) of the *T*. *zhukovskyi* subgenomes was markedly lower than that of domesticated timopheev’s wheat and einkorn wheat at the subgenomes level (Supplementary Table 2). We counted chromosome numbers of these plants and found only half showed the expected 42 chromosomes, while the remaining plants had abnormal chromosome counts, including plants having 41 (3 plants), 43 (5 plants), and 45 (2 plants) chromosomes, confirming karyotype instability (Source Data file, Supplementary Fig. 3).

We used normalized read counts to assess haplotype dosage of the subgenomes across the *T. zhukovskyi* chromosomes. A 2:2 homoeologous ratio with two A^m^ and two A^t^ chromosome copies would be expected in an allopolyploid species without homoeologous recombination. Homoeologous recombination can produce alterations from the 2:2 homoeologous ratio, resulting in copy-number variation (1:3, 3:1), and presence-absence variation (4:0, 0:4)^13^. Mapping of einkorn wheat and *T. timopheevii* reads against a synthetic reference combining assemblies of a domesticated einkorn wheat and *T. timopheevii* confirmed that the A^m^ and A^t^ subgenomes are sufficiently diverged to allow unambiguous read mapping (Supplementary Fig. 4, 5). Across the 19 *T. zhukovskyi* plants, we observed a complex mosaic of different chromosome ratios and extensive homoeologous exchange. Each plant displayed a unique pattern and various numbers of homoeologous exchanges (A^m^/A^t^) indicating high plant-to-plant variation within an accession (Supplementary Table 3, Source Data file).

For example, chromosome 7A of TA11273_P4 (plant number 4 of accession TA11273) exhibits a ∼450 Mb pericentromeric region with a 1:3 ratio for homoeologous segments of chromosome 7A^m^ :7A^t^ (Fig 2a, b). The telomeric ends of this chromosome had a 0:4 ratio while the interstitial regions showed a 4:0 ratio for 7A^m^:7A^t^, respectively (Fig. 2a, b). The telomeric and interstitial regions of chromosome 7A in this plant are thus fixed for the timopheev’s wheat and einkorn segments, respectively, analogous to nulliplex / quadruplex genotypes in an autopolyploid. This chromosome pattern suggested pairing and homoeologous exchange between the 7A^m^ and 7A^t^ chromosomes. To confirm the observations obtained from *in silico* data, we developed chromosome-specific painting probes for chromosomes 7A^m^ and 7A^t^. The probes were highly specific when hybridized to mitotic metaphase chromosome spreads of einkorn and *T. timopheevii* (Extended Data Fig. 3a). Hybridizing the probes to TA11273_P4 chromosome spreads confirmed the homoeologous exchanges between the A^m^ and A^t^ genomes (Fig. 2c). *T. zhukovskyi* plant TA11274_P3 (plant 3 of accession TA11274) showed a different segmental dosage pattern for chromosome 7A compared to TA11273_P4 (Extended Data Fig. 3b), confirming plant-to-plant variation. In addition, the progeny of TA11274_P3 revealed a different chromosome segregation ratio compared to their parental plant (Extended Data Fig. 3c), indicating tetrasomic inheritance of the A^m^ and A^t^ chromosomes. In summary, we show that the two A genomes in *T. zhukovskyi* recombine. The resulting mosaic-like pattern of allo- and autopolyploid genomic segments has been referred to as segmental allopolyploidy^11,13,14^.

Our results reveal extensive homoeologous exchange between the A^m^ and A^t^ genomes in *T. zhukovskyi* despite the presence of the pairing homoeologous (*Ph1*) locus^15,16^ indicating that the A genomes are not divergent enough for *Ph1* to prevent pairing. It is unclear, however, if the two A genomes only pair rarely or if there is frequent pairing and recombination. To address this question, we analysed populations of *T. timococcum* (a synthetic *T. zhukovskyi*) that were created from artificial crosses between einkorn wheat and *T. timopheevii*^17^. These populations enabled us to assess the frequency of homoeologous exchanges across defined numbers of generations (rounds of inbreeding). We produced and analysed skim-sequencing data^18^ from 88 individuals representing F_3_, F_5_, and F_12_ generations. We observed homoeologous exchanges already in the F_3_ generation, with an increasing number of homoeologous exchanges in the F_5_ and F_12_ generations, confirming that the A^m^ and A^t^ chromosomes pair and recombine frequently in each generation (Source Data file). The average number of homoeologous exchanges across the 19 analysed *T. zhukovskyi* plants was 22.8 (Supplementary Table 3, Source Data file).

### Transcriptional dynamics in *T. zhukovskyi*

*T. zhukovskyi* likely originated through a recent polyploidization event, raising intriguing questions about genetic and genomic changes that occur following polyploid formation. In particular, we were interested in examining changes in gene expression levels, given the presence of an additional A subgenome in *T. zhukovskyi.* Specifically, we sought to determine whether the increased A subgenome dosage in *T. zhukovskyi* leads to additive A-to-G expression levels relative to *T. timopheevii,* for which we should observe a 2:1 A-to-G relative expression level, or whether expression levels are balanced, resulting in similar A-to-G ratios (1:1) between the two species. To address this, we compared the ratio of A-to-G gene expression levels between the tetraploid and hexaploid species. Using a set of 11,607 single-copy triads, we found significant differences in the A-to-G gene expression ratios between tetraploid *T. timopheevii* and the synthetic hexaploid *T. timococcum*. While the A-to-G expression ratios in *T. timopheevii* were balanced (Fig. 2d, Supplementary Fig. 6), the median A-to-G expression ratios in *T*.

*timococcum* ranged from 1.98-2 (Fig. 2d, Supplementary Fig. 6, Supplementary Table 4). A similar result was obtained for hexaploid *T. zhukovskyi*, where the median A-to-G ratios ranged from 1.88 to 2 (Fig. 2d, Supplementary Fig. 7, Supplementary Table 5). For both *T. timococcum* and *T. zhukovskyi*, the overall results thus suggest that the hexaploid A-to-G expression ratios are at approximately double compared to the tetraploids, indicating additive effects of the additional A subgenome.

### *T. timopheevii* population genomics

In modern times, *T. timopheevii* cultivation has only been reported in a small geographical area in western Georgia^2^. This could indicate (i) that domesticated timopheev’s wheat represents a localized domestication event that resulted in an endemic wheat species with a very narrow distribution range, or (ii) that the domesticated *T. timopheevii* cultivated in Georgia might be the remains of a wheat species that had a much wider distribution range in historic times^2^. Recent archaeological studies reported that ‘new glume wheat’, excavated from archaeological sites across Europe, Anatolia and Transcaucasia, represent *T. timopheevii,* indicating that this wheat species might have been cultivated on a large scale until about 2,000 years ago^7,19,20^. To investigate the population genomics and evolutionary history of *T. timopheevii*, we generated whole-genome sequencing data (∼13.5-fold coverage) representing 178 wild and 47 domesticated accessions (Supplementary Table 6). The diversity panel has been selected to represent the geographical distribution of the species and to capture previously described population structures and diversity^21^.

In addition to domesticated *T. timopheevii*, a previous study identified two wild *T. timopheevii* populations named ARA-0 and ARA-1^2^. Our phylogenetic and ancestry analyses defined five *T. timopheevii* subpopulations (Fig. 3a, b, Extended Data Fig. 4 a-d, Supplementary Table 7). The domesticated *T. timopheevii* accessions formed a separate, narrow clade. The wild accessions grouped into four subpopulations, including ARA-1 and three subpopulations belonging to ARA-0 (Fig. 3a, b, Extended Data Fig. 4a, b, Supplementary Table 7). All ARA-1 accessions in our diversity panel were collected in a narrow geographical area spanning southern Türkiye and northwestern Syria. The subpopulations for ARA-0 correspond to ARA-0 NE comprising accessions from Azerbaijan, ARA-0 NW comprising accessions primarily collected in Armenia, and ARA-0 S including accessions from various regions in Iraq, Iran, and Türkiye.

**Fig. 3.**
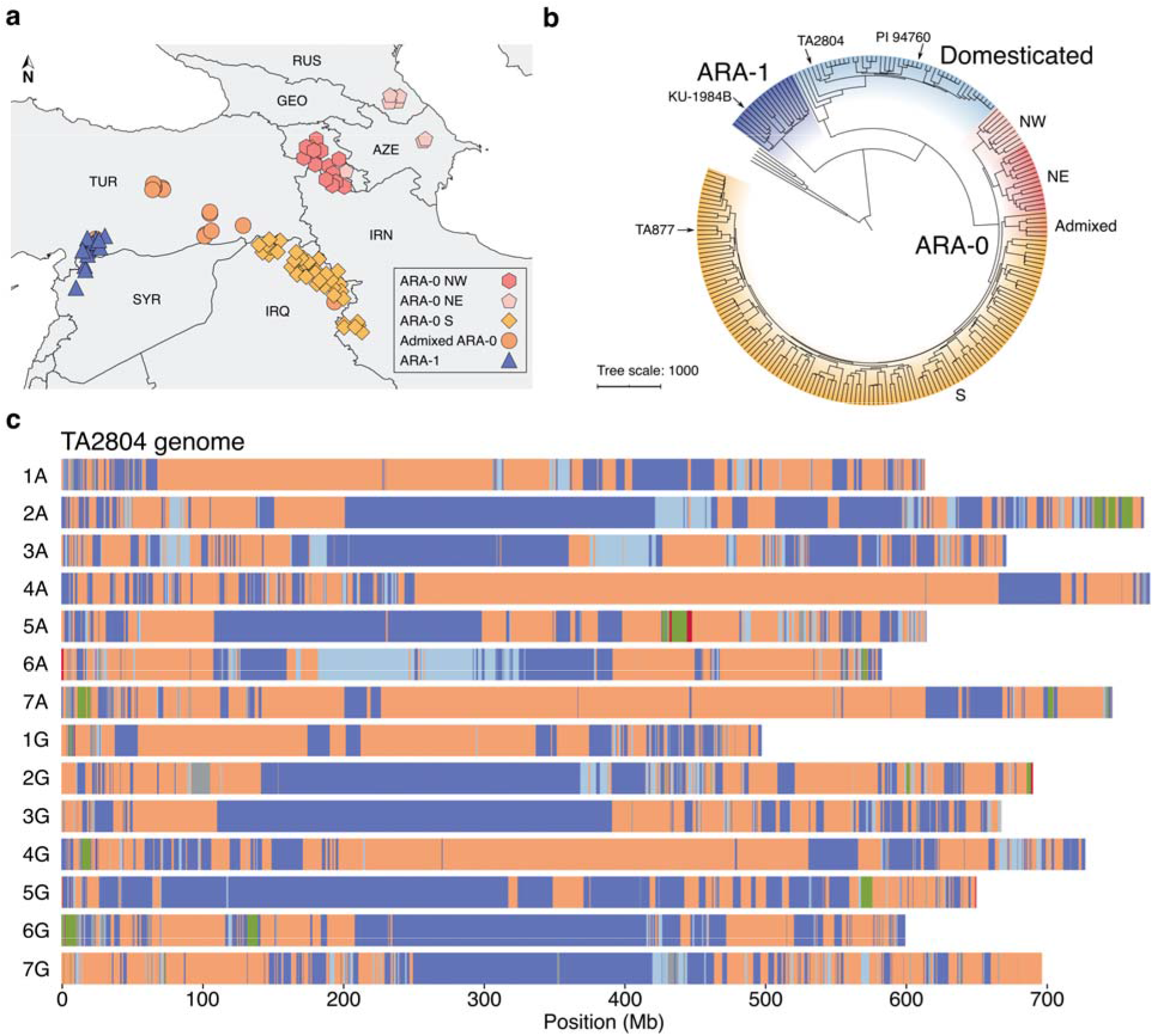
Population structure and domestication of *T. timopheevii*. **a,** Map showing the collection site and subpopulations of the wild *T. timopheevii* accessions. ARM, Armenia; AZE, Azerbaijan; GEO, Georgia; IRQ, Iraq; IRN, Islamic Republic of Iran; SYR, Syrian Arab Republic; TUR, Türkiye. **b,** SNP-based phylogenetic tree of *T. timopheevii* with five wild emmer wheat accessions as an outgroup (uncoloured branches). TA877 and TA2804 indicate the positions of the accessions for which chromosome-scale assemblies were generated in this study. A high-quality assembly for the domesticated *T. timopheevii* accession PI 94760 was generated in a previous study^6^. **c,** Haplotype composition of domesticated *T. timopheevii* accession TA2804. Haplotype segments being identical-by-state to ARA-0 are indicated in salmon colour and to ARA-1 in dark blue colour. Light blue colour shows *T. timopheevii* haplotype blocks that could not be assigned to ARA-0 or ARA-1. Haplotype segments shown in green colour originate from the Emmer wheat lineage and red colour from einkorn wheat. Unassigned windows are shown in grey colour.

The phylogenetic separation of domesticated *T. timopheevii* from the four wild subpopulations is intriguing. We found that domesticated *T. timopheevii* contains haplotype segments that are identical-by-state to all four wild subpopulations. In particular, domesticated *T. timopheevii* exhibits a mosaic-like haplotype pattern containing large chromosome segments originating from both ARA-0 and ARA-1, while these two main wild subpopulations remain genetically isolated with minimal evidence for gene flow between ARA-0 and ARA-1 accessions (Extended Data Fig. 5, 6). In total, around 38.4% (3.7 Gb) of the TA2804 genome is assignable to ARA-1, while around 49.9% (4.8 Gb) originated from ARA-0 (Fig. 3c). An additional 640 Mb (6.6%) of the TA2804 genome originates from *T. timopheevii* without being able to assign this portion of the genome to one of the two major wild populations. The remaining 5.1% (499 Mb) could not be assigned or were assigned to a non-*T. timopheevii* specific haplotype (Fig. 3c, Extended Data Fig. 6a). TreeMix analyses provides further support for gene flow from wild into domesticated *T. timopheevii* (Supplementary Fig. 8). These analyses establish the ARA-0 – ARA-1 admixture as a signature of domesticated *T. timopheevii*. Both the ARA-0 and ARA-1 populations contributed key domestication and adaptation genes to domesticated *T. timopheevii* (Supplementary Table 8). For the brittle rachis gene *Btr1-A*^22^ on chromosome 3A, all domesticated *T. timopheevii* accessions carry a 1 bp deletion resulting in a loss-of-function allele, while all wild accessions had the functional wild-type copy without the 1 bp deletion. For *Btr1-G* on chromosome 3G, ARA-1 accessions carry a functional copy, while ARA-0 and domesticated accessions carry a degenerated *Btr1-G* copy. The ARA-0 and ARA-1 accessions in our diversity panel only have a minimal geographical overlap today^2^. The patchwork-like genetic architecture of domesticated *T. timopheevii,* combining genomic segments from geographically isolated wild populations, is difficult to explain with a localized domestication event that resulted in an isolated domesticated *T. timopheevii* population restricted to Georgia. Consistent with archaeological findings, our results rather point to a wider cultivation range of domesticated *T. timopheevii* in the past, fostering gene flow from different wild subpopulations.

Consistent with a wider geographical distribution of domesticated timopheev’s wheat, we identified genomic signatures indicating gene flow from *T. timopheevii* into landraces from the Emmer wheat lineage that most likely occurred outside the current distribution ranges of wild and domesticated *T. timopheevii*. For example, we identified two independent but overlapping *T. timopheevii* introgressions on chromosome 5B in multiple bread wheat landraces (Fig. 4). The largest version of one of these introgressions, most likely reflecting its ancestral state, was most frequently found in bread wheat landraces collected from Afghanistan and India (Fig. 4). The two introgressions show admixture between ARA-0 and ARA-1, indicating an origin from domesticated *T. timopheevii.* Their exact haplotype compositions, however, do not match the ARA-0 - ARA-1 haplotype pattern present in the extant domesticated *T. timopheevii* gene pool, indicating that these introgressions originated from a different and now possibly extinct domesticated *T. timopheevii* population. In total, we identified 33 natural *T. timopheevii* introgressions with a cumulative size of 512 Mb across 1,943 wheat accessions representing the Emmer wheat lineage. Out of these 33 introgressions, 11 were identified as having a high likelihood of originating from extinct or unsampled domesticated *T. timopheevii* (Extended Data Fig. 7, Supplementary Table 9, 10). The *T. timopheevii* introgression segments in the Emmer wheat lineage contained a total of 11,786 gene models. Of these, 8,180 gene models were found in landraces with a maximum of 2,793 found in a single landrace and are most likely the result of natural gene flow from the Timopheevii into the Emmer wheat lineage (Supplementary Table 11).

**Fig. 4.**
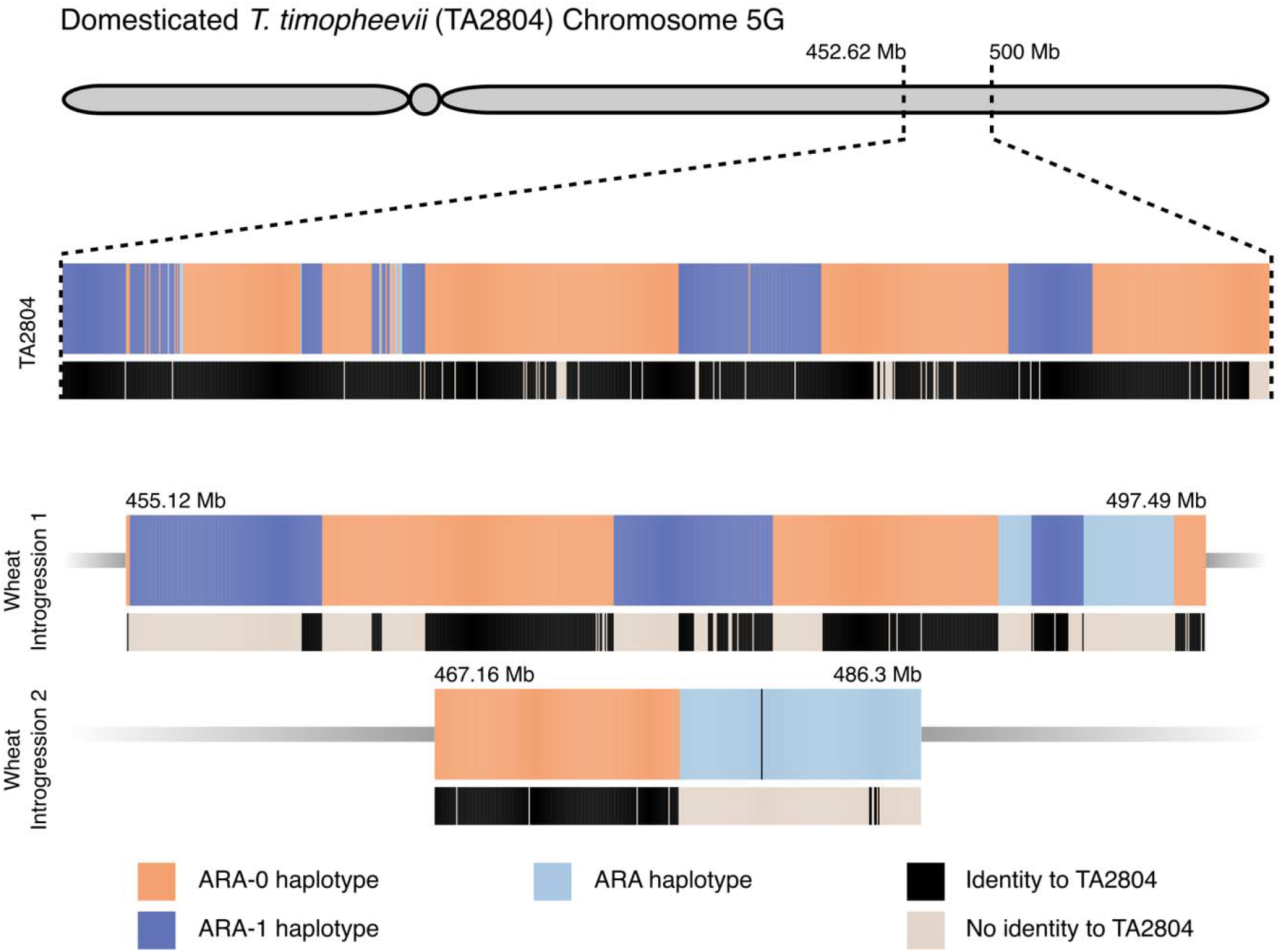
*T. timopheevii* introgressions into Emmer wheat lineage. Shown are schematic representation of three versions corresponding to syntenic *T. timopheevii* chromosome segments. The topmost version corresponds to the haplotype composition found on chromosome 5G of the domesticated *T. timopheevii* accession TA2804, with salmon colour indicating identity-by-state to ARA-0 and dark blue colour showing identity-by-state to ARA-1. Black bars in the track below the haplotype representation indicate that all the domesticated *T. timopheevii* and *T. zhukovskyi* accessions sequenced in our diversity panel showed identity-by-state to the TA2804 haplotype segment across the 50-Mb region, reflecting the restricted genetic diversity across extant domesticated *T. timopheevii*. The lower two versions correspond to *T. timopheevii* introgressions on chromosome 5B of wheat accessions representing the Emmer lineage. The larger version (introgression 1) is found in 25% of the landraces analysed and in 50.2% of th modern cultivars. The largest version of introgression 2 is found in landraces collected in Afghanistan and India, which is outside the current distribution range of domesticated and wild *T. timopheevii.* The grey bars in the lower tracks indicate chromosome segments that are not identical-by-state to the extant domesticated *T. timopheevii* and *T. zhukovskyi* gene pool, including TA2804, most likely indicating historic introgression from now-extinct domesticated *T. timopheevii* populations.

Together, these data indicate extensive gene flow between *T. timopheevii* and the Emmer wheat lineage, which is most likely the result of co-cultivation of timopheev’s wheat and wheat representing the Emmer lineage over a wide geographical range. Although *T. timopheevii* is no longer of global significance as a crop species, our data indicate that a significant proportion of genomic segments and genes in bread wheat are from *T. timopheevii*.

## Discussion

This study provides new insights into the genome composition, stability, and evolutionary dynamics of the Timopheevii wheat lineage. The mosaic-like haplotype pattern observed in domesticated *T. timopheevii* is similar to what has been seen in the Emmer lineage^23^ and also other crops^10,24–27^, indicating that crop domestication rarely follows a simple linear trajectory from a wild ancestor to a domesticated form^28^. Instead, recurrent gene flow from multiple wild populations have shaped domesticated crop genomes, resulting in patchwork-like haplotype compositions. In the case of domesticated *T. timopheevii*, contributions from ARA-0 and ARA-1 have resulted in its intermediate positioning within the phylogenetic tree.

Although *T. timopheevii* is of limited significance for global wheat production today, we discovered that a sizeable proportion of *T. timopheevii* genes persist in the bread wheat gene pool. Notably, we identified *T. timopheevii* haplotype segments that are no longer found in the extant domesticated *T. timopheevii* gene pool but are still maintained in bread wheat. These ‘fossil’ genetic footprints suggest that *T. timopheevii* cultivation was once much wider distributed. Since the 20^th^ century, *T. timopheevii* cultivation has been reported only in a limited geographical area, restricted to a few villages in Western Georgia^2^.

This has led to the hypothesis that Georgian *T. timopheevii* originated from a localized domestication event, resulting in an endemic population. However, the mosaic-like haplotype pattern of domesticated *T. timopheevii* uncovered in this study contradicts this notion. Instead, the most plausible explanation for its hybrid origin is the historical cultivation of *T. timopheevii* across a broad geographical range, where it came into contact with both wild ARA-0 and ARA-1 populations, facilitating gene flow. Our results agree with archaeological evidence suggesting a wider historical distribution of *T. timopheevii* cultivation across Europe and Western Asia in the past. Reports of ‘new glume wheat’ – now identified as *T. timopheevii* – from multiple agricultural settlements indicate its widespread cultivation until approximately 2,000 years ago^7,19,20^. It has been proposed that the expansion of the Roman Empire, coupled with sociocultural shifts such as urbanization, drove an increased demand for high-yielding wheat cultivar^7^, contributing to the gradual decline of *T. timopheevii* cultivation. The accessions cultivated in Georgia during the 20^th^ century are therefore likely the remnants of a wheat species that was once widely cultivated until the end of the Bronze Age. This raises an intriguing question: Have humans domesticated and cultivated additional *Triticum* or *Aegilops* species in the past that were not preserved as regional crops and have since been lost?

Another intriguing question pertaining to the Timopheevii lineage is the limited success of hexaploid species, which contrasts the case of hexaploid bread wheat where the additional genome provides greater adaptability and performance over the tetraploid. In the Emmer wheat lineage, durum and bread wheat are exceptional polyploid species in that their homoeologous chromosomes rarely or never pair, resulting in diploid-like genetic behaviour and high genome stability^15^. Many natural polyploids and polyploid crop species, on the other hand, exhibit some degree of homoeologous chromosome pairing and homoeologous exchange^13,29^. The correct pairing of homologous chromosomes in polyploid wheat is genetically regulated by the *Ph1* locus^15^. Despite the presence of *Ph1* in the Timopheevii wheat lineage^1,16,30^, we observed extensive homoeologous exchange in hexaploid *T. zhukovskyi*, suggesting that the A^m^ and A^t^ genomes are too closely related to prevent homoeologous chromosome pairing. The genomic instability of *T. zhukovskyi* may account for its limited agronomic significance, particularly for modern breeding, as selecting stable lines with consistent performance would be nearly impossible. In addition to genome stability, the success of the Emmer wheat lineage might also be due to ecological and agronomic factors.

In contrast, the karyotype stability of the Emmer wheat lineage likely played a crucial role in its emergence as a globally dominant crop lineage^31^. The chromosome stability within the Emmer wheat lineage has facilitated natural and artificial introgressions of alien chromosome segments from diverse wheat species and wild wheat relatives into the durum and bread wheat gene pools^32^. This gene flow may have been essential for restoring and increasing genetic diversity after domestication bottlenecks, enabling durum and bread wheat to adapt to various climatic conditions^10^. In contrast, hexaploid oat has undergone frequent chromosome rearrangements, which has created breeding barriers and limited alien introgressions^33^. Notably, although *T. zhukovskyi* has been classified as a distinct species^1^, the high sequence similarity among the accessions analysed in this study suggests that the *T. zhukovskyi* accessions maintained in *ex situ* collections may originate from a single genotype^2^. Thus, it appears that genome stability as a true allopolyploid is critical for broad success and particularly important for modern breeding of polyploid crops.

## Material and methods

### Plant material

We selected two accessions of *T. timopheevii*: a domesticated accession (*T. timopheevii* ssp. *timopheevii*), TA2804, and a wild accession (*T. timopheevii* ssp. *armeniacum*), TA877. The wild accession was collected near Erbil, Iraq, 4 km northeast of Shaqlawah (latitude: 36.4167, longitude: 44.3667; 1,000 m elevation). The domesticated accession was collected from the former USSR, but no further collection details are available. Both accessions have been preserved and maintained through single-seed descent at the Wheat Genetics Resource Center (WGRC) at Kansas State University. *T. zhukovskyi* accessions TA11273 and TA11274 (WGRC) in this study correspond to PI 352552 and PI 352553 of the USDA germplasm collection, respectively. According to the USDA original plant inventory, these two accessions were received from the Federal Agricultural Research Station in Switzerland in 1969. The Federal Agricultural Research Station received the two accessions from the Vavilov Research Institute. According to the available information, PI 352553 corresponds to the original *T. zhukovskyi* accession that was deposited in the Vavilov gene bank as accession number K-43063 in 1959^12^. Different generations of the synthetic hexaploid *T. timococcum* (2*n* = 6*x* = 42, GGA^t^A^t^A^m^A^m^) along with their parental accessions were provided by the Martonvásár Cereal Genebank (MVGB) at HUN-REN ATK, Martonvásár, Hungary^17^. In this paper we chose to adhere to the classification of the *Triticum* genus established by Mac Key in 2005^1^.

### Sequencing, assembly, annotation, and comparative analysis of *T. timopheevii* accessions TA2804 and TA877

#### PacBio HiFi library preparation and sequencing

High molecular weight DNA was extracted from young leaves of *T. timopheevii* using the NucleoBond HMW DNA Kit (MACHEREY-NAGEL, GmbH&Co.KG, Germany) according to the manufacturer’s instructions, and quantified using a NanoDrop One Spectrophotometer (Termo Fisher Scientifc, USA) and Qubit^®^ 4.0 Fluorometer with Qubit dsDNA BR Reagent (Termo Fisher Scientifc/Invitrogen, USA). DNA was sent to the Arizona Genomics Institute (AGI) for PacBio HiFi library preparation and sequencing of 10 SMRT cells for each *T. timopheevii* accession. Read quality was checked using FastQC (v0.11.8) (https://github.com/s-andrews/FastQC), adapters trimmed using fastp (v0.23.2)^34^ and reads were assembled into contig-level assemblies using hifiasm (v0.16)^35^ with default parameters.

For KU-1984B, DNA extraction, library preparation and the sequencing of four Revio SMRT cells was performed by the Arizona Genomics Institute (AGI) as a service, the resulting reads were assembled with hifiasm (v0.25.0)^35^ with default parameters.

#### Bionano optical map construction

To construct optical maps for the wild (TA877) and domesticated (TA2804) *T*. *timopheevii* accessions, we followed the procedure described in our previous study^9^, using the same protocols for high molecular weight (HMW) DNA extraction, labelling, and staining. Optical maps were then generated with the Bionano Genomics Saphyr System (Saphyr Chip G1.2) in accordance with the Saphyr System User Guide (3024-Bionano Genomics). Hybrid assemblies were generated using Bionano Solve (v3.7) software (https://bionanogenomics.com/support/software-downloads).

#### Generation of Hi-C reads

To generate Hi-C libraries for the two *T. timopheevii* accessions, cross-linked leaf samples were sent to Phase Genomics (Phase Genomics, Inc.). The libraries were prepared using restriction enzymes DPNII (GATC), DdeI (CTNAG), HinfI (GANTC), and MseI (TTAA), quality checked, and then sequenced on Illumina NovaSeq X+ by Psomagen Inc. (Rockville, Maryland, United States). For Hi-C scaffolding, the same strategy described in previous studies^9,36^ was used to achieve a high-quality, chromosome-scale assembly. First, the hybrid assemblies, which incorporated HiFi contigs and optical mapping data, were indexed using BWA (v0.7.17)^37^. Juicer (v1.6)^38^ was then used to align the Hi-C reads to the hybrid assemblies with default parameters with 64 threads to run, generating initial interaction matrices.

Following this alignment, 3D-DNA (v180419)^39^ was utilized to scaffold the assemblies, producing a draft Hi-C contact map for further refinement. Juicebox (v2.20.00)^40^ was used for visual inspection and manual adjustments to correct ordering and orientation of the pseudomolecules. To validate and further refine the assembly including orientation and ordering, Chromeister (v1.5.a)^41^ was used to align the draft assembly against a combined reference genome of *T. urartu* and *Ae. speltoides*. Finally, 3D-DNA (v180419)^39^ was re-run post-review to generate a finalized Hi-C contact map, completing the scaffolding process (Supplementary Fig. 1). One Hi-C library was generated and sequenced by the Arizona Genomics Institute (AGI) as a service for the scaffolding of KU-1984B. The reads were mapped with BWA mem (v0.7.17)^37^ following the Arima Genomic mapping pipeline’s default settings (https://github.com/ArimaGenomics/mapping_pipeline). The results of the mapping were then processed using YaHS (v1.1)^42^, followed by a few rounds of manual curation supported by Juicebox (v2.20.00)^40^ and Chromeister.

#### Transcriptome sequencing of T. timopheevii

RNA-seq data were generated from 20 tissues of TA877 and 18 tissues of TA2804, with RNA extraction (Supplementary Table 12), library preparation and sequencing conducted by Psomagen, Inc. generating target of ≥40 million reads per sample. For full-length isoform sequencing (Iso-seq), all tissue samples were pooled into two libraries per accession, prepared at AGI, and sequenced on three PacBio Sequel II SMRT cells per accession (two libraries per cell).

#### Gene prediction and transposable elements (TE) annotation

We performed *de novo* structural gene prediction, confidence classification, and functional annotation, following the protocol described by Mascher *et al*.^8^ using all the generated RNA-seq and Iso-seq data from wild (20 different tissues) and the domesticated TA2804 (18 different tissues) for gene prediction. Gene models for accession KU-1984B were constructed as consensus models derived from the annotations of TA877 and TA2804 using a gene projection approach that had been previously applied and described in a pan-barley genome study^43^. Detailed description of the methodology, parameterization, and implementation is available in the corresponding GitHub repositories (https://github.com/GeorgHaberer/gene_projection/blob/main/README.md and https://github.com/GeorgHaberer/gene_projection/tree/main/panhordeum). Differently from the pan-barley study, orthologous groups were inferred for *Brachypodium distachyon* (v3.1)^44^, barley (accession Morex), evidence-based annotations of the A, B, and D subgenomes of ten bread wheat genotypes^45,46^, as well as the A and G subgenomes of the two *T. timopheevii* lines of this study. All high-confidence, evidence-supported gene models of the *T. timopheevii* lines were used as source models for the projection. All remaining projection parameters were applied as previously described for the pan-barley pangenome. TE annotation was completed using EDTA^47^ with standard parameters. For further analysis the masking percentages of repetitive elements ranged from 76.4% to 84.5% (Supplementary Table 13-15).

#### Assembly quality assessment and validation

We evaluated assembly quality using multiple metrics. Contiguity statistics such as assembly length, number of contigs and contig N50 were assessed with QUAST (v 5.2.0)^48^. Completeness was determined using BUSCO (v5.7.1)^49^ with the poales_odb10 lineage dataset in euk_genome_min mode and miniport as gene predictor. Additionally, Merqury (v1.3)^50^ was used to estimate consensus quality (QV) and assembly completeness based on *k*-mer analysis using PacBio HiFi reads for *T. timopheevii* and DeepConsensus-corrected HiFi reads for *T. zhukovskyi*. To validate the *T. timopheevii* assemblies, we mapped PacBio HiFi reads using minimap2 (v2.24)^51^ and Illumina short reads using HISAT2 (v2.2.1)^52^ to the final assemblies checking total read mapping and non-reference variants (Extended Data Table 1). Variant calling using Illumina reads from TA2804 and TA877 to their respective assemblies showed negligible polymorphism, supporting assembly accuracy (Extended Data Fig. 8a, Supplementary Tables 16-18). The three assemblies synteny was confirmed by comparisons of gene models using GENESPACE (v1.3.1)^53^.

#### Chromatin immunoprecipitation (ChIP) and sequencing (ChIP-Seq) for TA2804

Chromatin immunoprecipitation (ChIP) for CENH3 was performed for TA2804 genome according to the method described in Ahmed *et al*. (2023)^9^. ChIP-seq library was constructed using the TruSeq ChIP Sample Prep Kit (Illumina, CA) according to the manufacturer’s instructions, and the libraries were sequenced using a NovaSeq X instrument with 2×150bp sequencing run at 4.8x genome coverage. The raw ChIP-Seq FASTQ files were trimmed using fastp (v0.23.2)^34^, and reads were aligned to the TA2804 assembly using HISAT2 (v2.2.1)^52^. Uniquely mapped concordant reads were extracted and used to compute coverage, using 10 kb bins to accurately identify centromere positions. As both biological replicates yielded highly consistent results, final peak calling and visualization was done using the combined data from both replicates, and functional centromere positions identified as a single peak on each chromosome (Supplementary Table 19). The custom scripts and BED files used for this analysis are available on GitHub. (https://github.com/laxmangene7/Timopheevii-Lineage-Genomics). DeepSeek (https://www.deepseek.com) was used to assist in annotating and refining the scripts.

#### Comparative analysis of wild and domesticated T. timopheevii

A Circos plot was generated to illustrate structural and functional genomic differences between the TA877, TA2804 and KU-1984B assemblies. The Circos was generated using the R (v4.3.0) package ‘circlize’ (v0.1.17)^54^, with feature density computed with a 10 Mb non-overlapping window, unless otherwise stated. OpenAI’s ChatGPT (https://chat.openai.com) was used to assist in generating scripts for Circos plotting. To dissect the synteny and orthology across the wild and domesticated *T. timopheevii* assemblies, we ran the R package GENESPACE (v1.3.1)^53^ to generate a comprehensive syntenic map between the assemblies. OrthoFinder (v2.5.5)^55^ was used to identify orthologous gene families across the assemblies with comparison of the high-confidence gene models and orthologs for collinearity between the two *T. timopheevii* assemblies.

### Contig-level assembly of *T. zhukovskyi*

Two individual plants of *T. zhukovskyi* accessions TA11273 and TA11274, named TA11273_P4 (plant #4) and TA11274_P3 (plant #3) respectively, were selected for PacBio HiFi sequencing based on (1) a chromosome number of 42, and (2) the most contrasting haplotype pattern across 19 *T. zhukovskyi* plants for which whole-genome sequencing data were generated. High-molecular weight (HMW) genomic DNA was extracted from young leaves of 3-week-old seedlings following a 48-h dark treatment grown at Kansas State University. Tissue was sent to AGI for HMW DNA extraction. The PacBio HiFi sequencing of the two *T*. *zhukovskyi* plants was done at Novogene (Novogene Inc.). Nine SMRT cells were produced for each plant and the raw subreads were pre-processed using DeepConsensus (v1.2.0)^56^ which resulted in 267 Gb (mean QV of 30) and 261 Gb (mean QV of 30.8) of HiFi data for TA11273_P4 and TA11274_P3, respectively. The corrected HiFi reads were assembled into contig-level assemblies using hifiasm (v0.19.8-r603)^35^ with default parameters. IBSpy (v0.4.6) (https://github.com/Uauy-Lab/IBSpy) was run with the contig-level assemblies of TA11273_P4 and TA11274_P3 as references and the domesticated einkorn (TA10622) and *T. timopheevii* (TA2804) as queries with a *k*-mer size of 31 and a window size of 50,000 bp as parameters. Windows were assigned based on the lowest variation score found in each window.

### Cytogenetics Analyses

#### Chromosome preparation and counting

Mitotic chromosome spreads were prepared from the root tips sampled from actively growing *T. zhukovskyi* and *T. timopheevii* plants following method described in Koo *et al*. (2018)^57^. The coverslips were removed with a double-edged razor blade, then slides were pre-treated with 20 μg/ml pepsin in 10 mM HCl for 3 min at 37 °C. Subsequently, slides were placed in distilled water for 1 min and they were washed three times with 2x SSC (300 mM Na-citrate, 30 mM NaCl, pH 7.0) for 5 min. The slides were treated with 4% formaldehyde in 2x SSC for 5 min, and dehydrated in ethanol series for 2 min each (70%, 90%, and 100%), and air-dried. Chromosome numbers were determined by analysing at least ten cells from each of two independent preparations in a plant.

#### Genomic in situ hybridization (GISH) and fluorescence in situ hybridization (FISH)

Genomic DNA of *T. timopheevii* was used as probes. For total genomic DNA, extraction of DNA from *T. monococcum* and *Ae. speltoides* and GISH procedure were as described by Koo *et al*. (2020)^58^. The procedures for mitotic chromosome preparation and fluorescence *in situ* hybridization (FISH) were adapted from Koo *et al*.^57^. Chromosome numbers were determined by analysing at least ten cells from each of two independent preparations in a plant. Haplotype-specific oligo probes^59^ for the chromosome 7A of einkorn accession TA10622 and chromosome 7A of *T. timopheevii* accession TA2804 were designed by Arbor Biosciences. Probes targeted Presence/Absence Variants (PAVs), SNPs, and indels, with 94,313 and 95,859 oligos synthesized for chromosome 7A of TA10622 and chromosome 7A of TA2804, respectively. Detailed information on the designed probes is available in the files provided in the Dryad repository. Each set was labelled with digoxigenin-11-dUTP or biotin-16-dUTP. The hybridization mixture (20 µl) contained 300 ng of each probe, 50% formamide, 2× SSC, and 10% dextran sulfate. Slides were denatured, hybridized for 18 h at 37 °C, and detected, captured and processed as in Koo *et al*. (2020)^58^.

### Skim-sequencing of T. zhukovskyi and T. timococcum

The skim-sequencing of *T. zhukovskyi* and *T. timococcum* used in this project followed the protocol of Nextera library preparation with slight modifications as described previously for skim-seq^18^. The pooled libraries were sequenced on Illumina NovoSeq X+ by Psomagen.

### Normalized read coverage analyses in *T. zhukovskyi* and *T. timococcum*

Each FASTQ file was trimmed using fastp (v0.23.2)^34^ using default parameters. The trimmed reads were then mapped using HISAT2 (v2.2.1)^52^ against an *in silico* synthetic reference which was a concatenation of the assemblies for domesticated einkorn accession TA10622^9^ and *T. timopheevii* TA2804. A custom script was used to calculate the number of uniquely mapped concordant reads mapped per 1-Mb bin (https://github.com/laxmangene7/Timopheevii-Lineage-Genomics). Read counts were normalized to genome coverage following the procedure described by Adhikari *et al*.^18^. The normalized read count reflects variation in chromosomal copy numbers, with values ranging from null (0 copy with normalized read depth ≍ 0) to four copies (normalized read depth ≈ 2). The normalized read counts were evaluated visually based on the coverage plots (Source Data file). The homoeologous exchanges between A^m^ and A^t^ subgenomes of *T*. *zhukovskyi* were calculated based on visual inspection of the coverage plots (Supplementary Table 3)

### Mapping accuracy estimation

Whole-genome sequencing data (∼5x coverage) from *T. zhukovskyi* plants, together with 50 domesticated einkorn and 47 domesticated *T. timopheevii* outgroup accessions, were mapped to a synthetic reference constructed by combining the assemblies of domesticated einkorn (TA10622)^9^ and *T. timopheevii* (TA2804). About 80% concordant reads were mapped uniquely using HISAT2 (v2.2.1)^52^ with a mis-mapping rate of < 0.2% (Supplementary Fig. 4). Mapping accuracy was independently validated by quantifying TA10622 and TA2804 reads assigned to the A^m^ and A^t^ subgenomes (Supplementary Fig. 5). We performed allele-matching^60^ analyses using A^m^-subgenome sites shared between *T*. *zhukovskyi* and *T*. *monococcum*, and A^t^ and G subgenome sites shared between *T*. *zhukovskyi* and *T*. *timopheevii*. These comparisons revealed that *T*. *zhukovskyi* individuals exhibit >98.5% allele matching, with the remaining ∼1.5% likely attributable to sequencing errors (Extended Data Fig. 2a, b).

### Gene expression analysis in *T. zhukovskyi* and *T. timococcum*

#### RNA extraction and sequencing

Three individual plants were grown for the accessions TA10622 (*T. monococcum*), TA2804 (*T. timopheevii*), and progeny of TA11273_P4 and TA11274_P3 (*T. zhukovskyi*). The samples were collected at the 2-leaf stage and total RNA extraction was performed using TRIzol™ reagent (Cat# 5596018; Invitrogen, USA). Approximately 100 mg of leaf tissue was flash-frozen in liquid nitrogen and ground to a fine powder using a pre-chilled mortar and pestle. The powdered tissue was homogenized in 1 ml of TRIzol reagent and incubated at room temperature for 5 min to ensure complete dissociation of nucleoprotein complexes. A volume of 200 µl of chloroform was added to the homogenate, followed by vigorous shaking and a 3-5 min incubation at room temperature. The samples were then centrifuged at 12,000 g for 10 min at 4 °C. The upper aqueous phase was carefully transferred to a new RNase-free tube, and RNA was precipitated by adding an equal volume of isopropanol. After a second centrifugation at 12,000 g for 10 min at 4 °C, the resulting RNA pellet was washed with 75% ethanol, briefly air-dried, and resuspended in RNase-free water. RNA integrity and quality were assessed by electrophoresis on a 1.5% agarose gel. Similarly, three individuals from the F12 generation of MVGB845-1T1 (*T. timococcum*) and its parents 1T-1 (*T. monococcum*) and MVGB845 (*T. timopheevii*) were grown in the KAUST greenhouse. Approximately 100 mg of frozen and ground tissue at the 2-leaf stage was used for RNA isolation using the Maxwell RSC Plant RNA Kit (AS1500) with a Maxwell RSC48 instrument following the manufacturer’s protocol (Promega). The Directional mRNA library preparation (poly A enrichment) and RNA sequencing were performed at Novogene using a target of 90 million paired-end reads for each sample.

#### Identification of single-copy triads

To investigate how *T. zhukovskyi* deals with the additional A subgenome dosage, we considered the A^t^ subgenome from *T. timopheevii* and the A^m^ subgenome from *T. monococcum* simply as AA subgenomes. A comparative gene dosage analysis was first carried out for *T. timococcum* and its known parental lines, *T. timopheevii* accession MVGB845 and *T. monococcum* accession 1T-1. For the comparison of subgenome dosage effects, first triads were identified. Triads were defined as genes present in all three (sub-)genomes in a single copy. Likewise, for the tetraploid analysis, dyads were defined as genes present in both sub-genomes in a single copy. For the identification of triads, the canonical protein sequences of *T. monococcum* TA10622 (A^m^ genome), *T. timopheevii* TA2804 (A^t^ subgenome) and *T. timopheevii* TA2804 (G subgenome) were selected and used as an input for Orthofinder (v2.5.5)^55^, which was executed with default parameters to identify orthologous genes.

#### Gene expression analyses

For gene expression analysis, the longest isoforms of high-confidence transcripts from *T. timopheevii* accession TA2804 and *T. monococcum* accession TA10622 were extracted with AGAT (v.0.8.0) (https://github.com/NBISweden/AGAT). A Kallisto (v.0.46.1)^61^ index of TA2804 was created and RNA-seq libraries from *T. timopheevii* (MVGB845 / 3 biological replicates) and *T. timopheevii* (TA2804 / 3 biological replicates) were aligned against the indexed transcriptome with Kallisto quant. The cDNA sequences of TA2804 and TA10622 were concatenated in order to obtain an *in silico* synthetic *T. zhukovskyi* transcriptome. Another Kallisto index for the synthetic transcriptome was generated. The RNA-seq libraries from two *T. zhukovskyi* (TA11273_P4 and TA11274_P3) with 3 biological replicates and a *T. timococcum* (an F12 plant) with 3 biological replicates were aligned to the indexed synthetic transcriptome with Kallisto quant. The resulting transcripts per million (TPM) values of the replicates for each dataset were used for pairwise comparison.

### Gene dosage analyses

For gene dosage analysis, the previously identified triads were used. Kruskal-Wallis and Dunn’s post-hoc tests were performed to determine whether the expression levels between different subgenomes differed significantly. To calculate the A-to-G ratios for both the hexaploid verses tetraploid species, we focused on triads with TPM values for A^m^ + A^t^ > 0.5 and G > 0.5 and on dyads with TPM values for A^t^ > 0.5 and G > 0.5, respectively. The A-to-G ratio for each subset of triads and dyads was calculated and the resulting value for the triad was divided by that of the corresponding dyad based on orthology information. This resulted in a set of 7,022 to 7,703 homeologs (Supplementary Tables 4, 5). A total of 9 pairwise comparisons between each of the biological replicates of *T. timopheevii* MVG845 and *T. timococcum* replicates was carried out. For *T. timopheevii* TA2804 and *T. zhukovskyi*, a total of 18 pairwise comparisons were conducted. OpenAI’s ChatGPT (https://chat.openai.com) was used to assist in refining and debugging the Python script used for the gene dosage analyses.

### *T. timopheevii* population genomics

#### Population selection and sequencing

We compiled a diversity panel comprising 233 *T. timopheevii* accessions (Supplementary Table 6), representing 178 wild (*T. timopheevii* ssp*. armeniacum)* and 47 domesticated (*T. timopheevii* ssp*. timopheevii*) accessions. One accession (TRI 7258) was identified as misclassified based on its position in the phylogenetic tree and was subsequently removed. The diversity panel was selected to maximize the geographical representation and the translocated karyotypes previously reported in the wild population^2^. From the 76 variants of chromosomal rearrangements described, we covered 50 different translocation types. The diversity panel was grown and sequenced in two batches; 142 accessions were grown as single plants in the KSU greenhouse, and 91 in the KAUST greenhouse keeping similar environmental condition. Genomic DNA was extracted from one or two young leaves of a single plant. Leaf tissue, approximately 1–1.5 cm in size, was harvested three weeks after germination and lyophilized for three days followed by grinding using a Geno/Grinder at 1,400 rpm for two minutes. DNA extraction was carried out using the MagMax extraction kit, with purification conducted on the KingFisher Flex system. Library preparation, quality control (QC) and sequencing were performed by Psomagen (Psomagen, Inc.). TruSeq DNA PCR-free (350) libraries were generated, and fragment size and QC were checked. Sequencing was performed on the NovaSeq X platform and 150 bp paired-end reads were obtained.

#### Read mapping and SNP calling

The raw FASTQ files were trimmed using fastp (v0.23.2)^34^ with a Phred quality score threshold of 20 and a minimum read length of 150 bp. The trimmed high-quality reads were aligned using HISAT2 (v2.2.1) aligner^52^ and variants were called on both the domesticated TA2804 and the wild TA877 reference assemblies separately. Following alignment using SAMtools/BCFtools (v1.16)^62^, the resulting SAM files were converted to BAM format, and the files were sorted and indexed and used for variant calling. To remove low-quality sites, we filtered the variants in two steps: i) using BCFtools filter parameters to keep the SNPs with QUAL ≥ 30, INFO/DP ≥ 20, F_MISSING ≤ 0.2, AF > 0.01, and AF < 0.99; ii), keep SNPs with MAF ≥ 0.01 and heterozygosity < 10% using a custom script. The first BCFtools pass applied basic quality thresholds while it could also retain multiallelic variants (AF > 0.01). The second filter enforced stricter criteria, focusing on segregating biallelic high quality SNPs (MAF ≥ 0.01, heterozygosity ≤ 0.10) (Supplementary Table 16). For computational efficiency in subsequent analyses and preservation of representative genomic information, we used the ’SelectVariants’ option in GATK (v4.3.0.0)^63^, to select a 10% subset of the filtered variants from each of the 14 chromosomes. The merged VCF file was converted to hapmap file using tassel (v5.2.60)^64^. Variant calling with whole-genome sequencing data of the reference accessions TA877 and TA2804 mapped to their respective reference assemblies, was used to assess accuracy of the genotyping by calculating the percentage of polymorphic loci relative to the total non-missing sites. The polymorphic loci percentages were 0.00074% for TA2804 (Supplementary Table 17) and 0.00081% for TA877 (Supplementary Table 18), underscoring the high quality of the assemblies, read alignment, and SNP filtering. The genome-wide SNP density revealed a non-uniform distribution of SNPs across the genome, with higher SNP density in telomeric regions compared to pericentromeric regions (Extended Data Fig. 8b, 9a).

### Population structure and diversity

We explored the population structure of *T. timopheevii* using phylogeny and population structure. Phylogenetic clustering was performed by calculating genetic distances using the dist function in R (v4.3.0) with Euclidean method. The resulting distances were converted into a phylogenetic object using the ape (v5.8.1)^65^ R package, and the neighbour-joining (NJ) tree was visualized using the phyclust (v0.1-34)^66^ R package. In the unrooted NJ tree, branches were colour-coded according to their genetic grouping (Fig. 3b, Extended Data Fig. 9b).

Population structure analysis was carried out in R (v4.3.0) using the Landscape and Ecological Association Studies (LEA) package (v3.14.0)^67^. Least-squares estimates of ancestry proportions and ancestral allele frequencies were computed using the ‘snmf’ function, assuming *K*=5 ancestral populations. The optimal number of population subgroups (*K*) was determined based on cross-entropy values, reflecting the maximal local contribution of ancestry^9^ (Extended Data Fig 3b, Supplementary Table 7, 20, Extended Data Fig. 9c). The ancestry proportions, represented by the Q matrix, were visualized using the POPHELPER (v2.3.1)^68^ R package. To minimize the potential bias in subpopulation differentiation due to subgroup sizes, we selected a subset of the entire population, particularly by reducing the size of ARA-0 subgroup through random selection. The subset includes 44 ARA-0, 17 ARA-1 and 17 domesticated accessions (Extended Data Fig. 4c). The genetic diversity of the *T. timopheevii* population was assessed by computing Nei’s diversity indices^69^, nucleotide diversity (π), and the total number of segregating loci for specific population subsets. For both indices, subsets were extracted from the VCF file using VCFtools (v0.1.17)^70^. Nei’s index was calculated as previously described in Adhikari *et al*. (2023)^71^, while π was computed using a 1-Mb sliding window in the separated VCF file for the subset^9^, and the mean π values were derived accordingly (Supplementary Table 21). To determine the total number of segregating loci within subgroups, we removed sites with entirely missing genotypes or those that were completely fixed (MAF = 0) in the specific subgroup, then counted the remaining segregating loci for each group. We computed pairwise Fst (Weir and Cockerham’s Fst^72^) between the ARA-0, ARA-1 and domesticated populations using VCFtools (v0.1.17)^70^ on the down sampled dataset. The pairwise mean Fst was calculated in 100-kb bins for all three population pairs: ARA-0 vs. ARA-1, ARA-0 vs. domesticated, and ARA-1 vs. domesticated. Subsequently, a genome-wide mean Fst was computed for each pair resulting in 0.5, 0.38 and 0.43 respectively.

A distinct filtering, similar to the one described in Guo *et al.* 2025^27^, was applied on the raw VCF file containing the variants called over the TA2804 reference. First, we used BCFtools (v1.16)^62^ to retain biallelic SNPs with the following quality settings: -q 20 -Q 20. Subsequently, all homozygous SNPs showing a depth lower than 2 and higher than 50 and heterozygous SNPs with the depths of both alleles not being greater or equal to three were set to ‘missing’.

We used PLINK (v 1.90b6.24)^73^ to assign the allele frequencies to the five identified subpopulations. We subsequently used TreeMix (v1.3)^74^ to infer migration events with the following parameters: -m 1 -k 1000 -global.

### k-mer generation

We generate 31-mer sets of the 225 re-sequenced *T. timopheevii* accessions, 617 accessions of publicly available wild wheat relatives and 1,943 publicly available wheat accessions (Supplementary Table 10, 22). The raw sequences were trimmed using Trimmomatic (v0.39)^75^ (LEADING:3 TRAILING:3 SLIDINGWINDOW:4:25 MINLEN:75) and the *k*-mers were counted with KMC3 (v3.1.2)^76^ retaining all *k*-mers.

### FastIBS

In order to improve the efficiency of the IBSpy pipeline (https://github.com/Uauy-Lab/IBSpy), we created a faster C based version able to use parallelization. The pipeline’s main function is ‘fastibs’ that produces the same results as the Identity-by-State Python pipeline^9^. Similar to IBSpy, FastIBS (https://github.com/githubcbrc/FastIBS) uses KMC3 *k*-mer databases as input. Unlike IBSpy, however, FastIBS can process multiple reference genomes at the same time.

### k-mer-based phylogeny

Considering the availability of 31-mers sets for the evolutional analysis, we tested a *k*-mer based phylogeny. Using the function ‘KDBIntersect’ of FastIBS (https://github.com/githubcbrc/FastIBS), we calculated the distance between each *T. timopheevii* sample sequenced in this study (Supplementary Table 6) by computing the pairwise number of common *k*-mers between all the *k*-mer sets. The distance between each of the sets was normalized with the following formula:

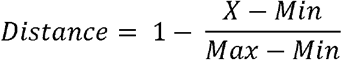

Where ‘X’ is the number of common *k*-mers between the two sets the distance is measured from, ‘Min’ is the minimal number of common *k*-mers found among all the comparisons and ‘Max’ is the maximum number of common *k*-mers found among all the comparisons. The normalized distance matrix (Supplementary Table 23) was converted into a distance network with the Python package NetworkX (v3.4.2)^77^ with the minimum threshold to set an edge between the nodes being 0.32 (Extended Data Fig. 4d).

### Genes introgressed in wheat gene pool

To achieve a complete analysis of *T. timopheevii*-specific genes present in wheat, the FastIBS (https://github.com/githubcbrc/FastIBS) function ‘fastibsmapper’ was used. The function uses a FASTA file and a *k*-mer set as inputs, and provides a csv file as output, in which for each field, corresponding to a base of the input FASTA file, sum of all the *k*-mers that can perfectly cover the base is reported. In case of 31-mers used in this study, a perfect identity is shown as a sequence of 31 as value. SNPs or deletions are visualized as valleys for which there is a gradual decrease in the *k*-mer coverage until values close to 0 are reached. The 88,243 TA2804 gene models were compared to 2,244 31-mer sets comprising 1,943 cultivated wheat accessions, 78 wild emmer accessions, and 178 wild *T. timopheevii* accessions using FastIBS fastibsmapper function. For each accession and for each gene model, the set of per-base numbers of mapped *k*-mers were summed up. For each gene model, we computed the theoretical maximum sum value based on the length of the gene model and assuming 100% identity between the accession and the reference ((length - 60)*31 + 465 *2). Each actual sum value calculated for each gene model was then subtracted from the theoretical maximum value, resulting in a distance score where 0 indicates 100% identity between the accession and the TA2804 allele. For each gene model, the average distance score for wild emmer, ARA-0, and ARA-1 were computed. For each gene model, the respective gene in each wheat cultivar or landrace of the Emmer wheat lineage were reported as potentially originating from *T. timopheevii* if at least one of the two following conditions was true: its distance from TA2804 was smaller than the distance between TA2804 and the average of ARA-0 plus a tolerance value, or its distance from TA2804 was smaller than the ARA-1 average distance plus a tolerance value. The tolerance value was calculated for each gene model as 10% of the difference between the wild emmer average distance subtracted from the ARA-0 average distance. The ARA-1 tolerance value was computed the same way by using the ARA-1 average distance instead of ARA-0 in the subtraction with wild emmer average distance. The three averages are reported for each gene model in the output file for quality control. For some genes it was not possible to compute the allele distance with enough certainty because of the average distribution. Gene models are retained ‘valid’ if both the average distances between TA2804 and ARA-0 and ARA-1 were greater than to wild emmer (to exclude genes that were introgressed from the Emmer wheat lineage into TA2804) and if the value resulting from the following formula is between 2/3 and 3/2: x = (AvWEW - AvARA-0)/(AvWEW AvARA-1). This was to ensure that the two ARA subpopulations are close together and far from wild emmer. An enrichment analysis was performed using GOATOOLS (v1.1.5)^78^, the study set was made from all genes found in landraces considered valid for which the average was in the range of ARA-0 or ARA-1 and the background set made using all valid genes. An enrichment for the GO terms corresponding to ‘response to auxin’, ‘electron transport coupled proton transport’, ‘defense response to fungus’ and ‘aerobic respiration’ was found significative. A depletion for the GO term corresponding to ‘RNA metabolic process’ was found significative.

### Determine subpopulation contributions to T. timopheevii

The analysis based on the FastIBS (https://github.com/githubcbrc/FastIBS) output to determine the origin of the haplotypes of the 50-kb windows of TA2804, TA877 and PI 94760 is described in Supplementary Note 1.

### Introgressions analysis

The complete workflow of the assessment of introgressions from *T. timopheevii* wheat into domesticated Emmer lineage wheat and the putative attribution to New Glume Wheat is described in Supplementary Note 2.

### Domestication genes analysis

We used BLAST to identify the sequences and positions of the following genes: *Pdp-1A*, *Pdp-1G*, *Vrn-1A*, *Vrn-1G*, *PhyC-A*, *PhyC-G*, *Btr-1A*, *Btr-1G*. We used FastIBS fastibsmapper with the gene sequences as reference using the *k*-mer sets for all the Timopheevii lineage accessions available in this study to infer the closest wild population carrying the same allele as the domesticated group.

### Map generation

The map in Fig. 3a was generated using QGIS (v3.40.3).

## Data availability

The raw Illumina reads files for the whole-genome resequencing panel of *T*. *timopheevii* used in this study were deposited at NCBI under BioProject number PRJNA1258594. The *T*. *timopheevii* multi-omics data including raw PacBio HiFi, Hi-C, Iso-Seq, RNA-Seq, and CENH3 ChIP-seq FASTQ files of accession TA2804 are available at NCBI under BioProject number PRJNA1262074. Corrected HiFi reads of two *T*. *zhukovskyi* accessions, whole-genome sequencing FASTQ files of *T*. *zhukovskyi* and *T*. *timococcum* parental lines (*T*. *timopheevii* and *T*. *monococcum*), RNA-Seq FASTQ files of *T*. *zhukovskyi* and *T*. *timococcum*, and 95 skim-seq data (88 *T*. *timococcum* lines, six *T*. *timopheevii* and one *T*. *monococcum*) are available under NCBI BioProject number PRJNA1263130. Two chromosome-scale genome assemblies of *T*. *timopheevii* (GCA_965283245 and GCA_965283015) and the contig-level genome assemblies of the two *T*. *zhukovskyi* accessions were deposited in the European Nucleotide Archive (ENA) under number PRJEB89274. The two genome assemblies of *T*. *zhukovskyi*, the two chromosome-scale genome assemblies of *T*. *timopheevii* and the annotation files *T*. *timopheevii*, the SNP matrices used in the diversity panel analysis, VCF files used for *T*. *timopheevii* population genomics, wheat chromosomes probe set detail, input files necessary for the gene dosage analysis and the resulting output files are available at Dryad (https://doi.org/10.5061/dryad.ksn02v7g7).

Publicly available wild wheat relative sequencing data were retrieved from NCBI under project numbers PRJNA759292, PRJNA337888, PRJNA628827, PRJNA476679, PRJNA310175; at EBI-ENA under project number PRJEB61155; at China National GeneBank under project number CNP0000699; at Genome Sequence Archive in the BIG Data Center under the project number PRJCA001626.

Publicly available Emmer lineage domesticated wheat sequencing data were retrieved from NCBI under project numbers PRJNA1070409, PRJNA1188632, PRJNA439156, PRJNA476679, PRJNA596843, PRJNA597250, PRJNA663409, PRJNA669381, PRJNA670578, PRJNA694980, PRJNA714281, PRJNA722149, PRJNA729723, PRJNA744310, PRJNA745496, PRJNA759292, PRJNA771357, PRJNA790490, PRJNA820989, PRJNA877303, PRJNA900700, PRJNA918327, PRJNA956839, PRJNA986484, PRJNA986532; at EBI-ENA under project numbers PRJEB22687, PRJEB44721, PRJEB45541, PRJEB48529, PRJEB49351; at Genome Sequence Archive in the BIG Data Center under the project numbers PRJCA004228, PRJCA004273, PRJCA005979, PRJCA009783, PRJCA019508, PRJCA019636, PRJCA021345.

## Code availability

The script developed for the gene-dosage analysis is available at https://github.com/samaralemos/gene_dosage_analysis. The scripts used to generate Circos plots, read mapping coverage graphs (including ChIP-seq and whole-genome sequencing in translocated regions), skim-seq data visualizations, copy number profiles, and the *T*. *zhukovskyi* heat maps are available at https://github.com/laxmangene7/Timopheevii-Lineage-Genomics. The scripts used to perform the haplotype composition analysis are available at https://github.com/emilecg/T.timopheevii_evolution. FastIBS is available at https://github.com/githubcbrc/FastIBS.

## Acknowledgements

This research used the Shaheen supercomputer and the Ibex cluster managed by the KAUST Supercomputing Laboratory (KSL) at King Abdullah University of Science and Technology (KAUST). We thank the KSL systems administrators and computational scientists for help with debugging and overall support. We are grateful to Hanan Sela of the University of Haifa for care of the *T*. *timopheevii* plants grown for assemblies, transcriptomics, and tissue collection. We thank Shuangye Wu and Ethan Faryna for preparing the skim-sequencing libraries and extracting DNA for a part of a *T. timopheevii* whole-genome resequencing panel. We also thank Priyanka Kalambettu, Gabrielle Michaux, Ibrahim Elbasyoni, and Xingfang Shi for their assistance with DNA extraction and care of the resequencing panel in the greenhouse and vernalization chambers. This publication is based upon work supported by KAUST award ORFS-CRG11-2022-5076 to S.G.K, baseline funding from KAUST to J.P. and S.G.K, and USDA National Institute of Food and Agriculture (NIFA) grant no. 2022-67013-36362 to J.P. A.S. and X.G. were supported by the King Abdullah University of Science and Technology (KAUST) Center of Excellence for Smart Health (KCSH), under award number 5932. This project was also supported by WGRC/IUCRC and NSF (grant #1822162) funded to D-H.K. This project was also supported by the US-Israel Binational Agricultural Research and Development Fund (BARD project No. IS-5188-19), and the Israel Science Foundation (ISF grant 1431/22 and 2223/22) funded to A.D.

## Author Contributions

M.A., L.A., A.D., J.P and S.G.K. conceived the project and designed the sequencing strategy. A.D., J.P. and S.G.K selected accessions for assembling. N.R. constructed the TA877 and TA2804 optical maps. M.A., L.A. and E.C.G. assembled the *T. timopheevii and T. zhukovskyi* genomes and performed quality checks. T.L., G.H. and M.S. performed gene model prediction on the *T. timopheevii* assemblies. M.A. and L.A. performed dosage analysis in *T. zhukovskyi*. D-H. K. performed karyotype analysis, ChIP-Seq and cytogenetic experiments. P.M. and I.M. generated the *T. timococcum* populations. S.C.L. and N.K. performed gene expression analysis. M.A. and H.Ö. configured *T. timopheevii* diversity panel. W.J.R. and H.Ö. maintained the diversity panel and distributed seed. A.S. and X.G. developed FastIBS. L.A. performed comparative genomics analysis between *T. timopheevii* assemblies, CENH3 ChIP-Seq mapping, and resequencing panel read mapping, SNP calling and quality control. L.A. and E.C.G. performed phylogenetic and ancestry analyses. E.C.G. performed haplotype composition analysis. E.C.G performed the introgression analysis. S.G.K., J.P., L.A., M.A. and E.C.G drafted the first version of manuscript. All authors read, commented on and approved the final version of the manuscript.

## Ethics declarations

Competing interests

The authors declare no competing interests.

## References

1. Feldman, M. & Levy, A.A. Wheat Evolution and Domestication, (Springer Nature, Switzeland, 2023).

2. Badaeva, E.D. et al. Genetic diversity, distribution and domestication history of the neglected GGA^t^A^t^ genepool of wheat. Theoretical and Applied Genetics 135, 755–776 (2022).

3. Xiong, W. et al. New wheat breeding paradigms for a warming climate. Nature Climate Change 14, 869–875 (2024).

4. Tadesse, W. et al. Genetic gains in wheat breeding and its role in feeding the world. *Crop Breeding*, Genetics and Genomics 1, e190005 (2019).

5. Kilian, B. et al. Independent wheat B and G genome origins in outcrossing *Aegilops* progenitor haplotypes. Molecular Biology and Evolution 24, 217–227 (2007).

6. Grewal, S. et al. Chromosome-scale genome assembly of bread wheat’s wild relative *Triticum timopheevii*. Scientific Data 11, 420 (2024).

7. Filipović, D. et al. *Triticum timopheevii* s.l. (‘new glume wheat’) finds in regions of southern and eastern Europe across space and time. Vegetation History and Archaeobotany 33, 195–208 (2024).

8. Mascher, M. et al. Long-read sequence assembly: a technical evaluation in barley. The Plant Cell 33, 1888–1906 (2021).

9. Ahmed, H.I. et al. Einkorn genomics sheds light on history of the oldest domesticated wheat. Nature 620, 830–838 (2023).

10. Cavalet-Giorsa, E. et al. Origin and evolution of the bread wheat D genome. Nature 633, 848–855 (2024).

11. Mason, A.S. & Wendel, J.F. Homoeologous exchanges, segmental allopolyploidy, and polyploid genome evolution. Frontiers in Genetics 11, 1014 (2020).

12. Badaeva, E.D. et al. Molecular cytogenetic characterization of *Triticum timopheevii* chromosomes provides new insight on genome evolution of *T. zhukovskyi*. Plant Systematics and Evolution 302, 943–956 (2016).

13. Deb, S.K., Edger, P.P., Pires, J.C. & McKain, M.R. Patterns, mechanisms, and consequences of homoeologous exchange in allopolyploid angiosperms: a genomic and epigenomic perspective. New Phytologist 238, 2284–2304 (2023).

14. Stebbins Jr, G.L. Types of polyploids: their classification and significance. Advances in Genetics 1, 403–429 (1947).

15. Griffiths, S. et al. Molecular characterization of *Ph1* as a major chromosome pairing locus in polyploid wheat. Nature 439, 749–752 (2006).

16. Ozkan, H. & Feldman, M. Genotypic variation in tetraploid wheat affecting homoeologous pairing in hybrids with *Aegilops peregrina*. Genome 44, 1000–1006 (2001).

17. Mikó, P., Megyeri, M., Farkas, A., Molnár, I. & Molnár-Láng, M. Molecular cytogenetic identification and phenotypic description of a new synthetic amphiploid, *Triticum timococcum* (A^t^A^t^GGA^m^A^m^). Genetic Resources and Crop Evolution 62, 55–66 (2015).

18. Adhikari, L. et al. A high-throughput skim-sequencing approach for genotyping, dosage estimation and identifying translocations. Scientific Reports 12, 17583 (2022).

19. Czajkowska, B.I. et al. Ancient DNA typing indicates that the “new” glume wheat of early Eurasian agriculture is a cultivated member of the *Triticum timopheevii* group. Journal of Archaeological Science 123, 105258 (2020).

20. Roushannafas, T., Bogaard, A. & Charles, M. Geometric morphometrics sheds new light on the identification and domestication status of ‘new glume wheat’ at Neolithic Çatalhöyük. Journal of Archaeological Science 142, 105599 (2022).

21. Yadav, I.S. et al. Exploring genetic diversity of wild and related tetraploid wheat species *Triticum turgidum* and *Triticum timopheevii*. Journal of Advanced Research 48, 47–60 (2023).

22. Nave, M. et al. The independent domestication of Timopheev’s wheat: insights from haplotype analysis of the *Brittle rachis 1* (*BTR1*-*A*) gene. Genes 12, 338 (2021).

23. Wang, Z. et al. Dispersed emergence and protracted domestication of polyploid wheat uncovered by mosaic ancestral haploblock inference. Nature Communications 13, 3891 (2022).

24. Kistler, L. et al. Historic manioc genomes illuminate maintenance of diversity under long-lived clonal cultivation. Science 387, eadq0018 (2025).

25. Yang, N. et al. Two teosintes made modern maize. Science 382, eadg8940 (2023).

26. He, F. et al. Exome sequencing highlights the role of wild-relative introgression in shaping the adaptive landscape of the wheat genome. Nature Genetics 51, 896–904 (2019).

27. Guo, Y. et al. A haplotype-based evolutionary history of barley domestication. Nature 647, 680–688 (2025).

28. Goncalves-Dias, J., Singh, A., Graf, C. & Stetter, M.G. Genetic incompatibilities and evolutionary rescue by wild relatives shaped grain amaranth domestication. Molecular Biology and Evolution 40, msad177 (2023).

29. Zhang, Z. et al. Homoeologous exchanges occur through intragenic recombination generating novel transcripts and proteins in wheat and other polyploids. Proceedings of the National Academy of Sciences of the United States of America 117, 14561–14571 (2020).

30. Martinez, M., Naranjo, T., Cuadrado, C. & Romero, C. Synaptic behaviour of the tetraploid wheat *Triticum timopheevii*. Theoretical and Applied Genetics 93, 1139–1144 (1996).

31. Zhang, H.K. et al. Intrinsic karyotype stability and gene copy number variations may have laid the foundation for tetraploid wheat formation. Proceedings of the National Academy of Sciences of the United States of America 110, 19466–19471 (2013).

32. Molnar-Lang, M., Ceoloni, C. & Dolezel, J. Alien Introgression in Wheat, (Springer Cham, 2015).

33. Kamal, N. et al. The mosaic oat genome gives insights into a uniquely healthy cereal crop. Nature 606, 113–119 (2022).

34. Chen, S. Ultrafast one-pass FASTQ data preprocessing, quality control, and deduplication using fastp. iMeta 2, e107 (2023).

35. Cheng, H., Concepcion, G.T., Feng, X., Zhang, H. & Li, H. Haplotype-resolved de novo assembly using phased assembly graphs with hifiasm. Nature Methods 18, 170–175 (2021).

36. Abrouk, M. et al. Chromosome-scale assembly of the wild wheat relative *Aegilops umbellulata*. Scientific Data 10, 739 (2023).

37. Li, H. Aligning sequence reads, clone sequences and assembly contigs with BWA-MEM. arXiv preprint arXiv:1303.3997 (2013).

38. Durand, N.C. et al. Juicer Provides a one-click system for analyzing loop-resolution Hi-C experiments. Cell Systems 3, 95–98 (2016).

39. Dudchenko, O. et al. De novo assembly of the *Aedes aegypti* genome using Hi-C yields chromosome-length scaffolds. Science 356, 92–95 (2017).

40. Durand, N.C. et al. Juicebox provides a visualization system for Hi-C contact maps with unlimited zoom. Cell Systems 3, 99–101 (2016).

41. Pérez-Wohlfeil, E., Diaz-del-Pino, S. & Trelles, O. Ultra-fast genome comparison for large-scale genomic experiments. Scientific Reports 9, 10274 (2019).

42. Zhou, C., McCarthy, S.A. & Durbin, R. YaHS: yet another Hi-C scaffolding tool. Bioinformatics 39, btac808 (2022).

43. Jayakodi, M. et al. Structural variation in the pangenome of wild and domesticated barley. Nature 636, 654–662 (2024).

44. Goodstein, D.M. et al. Phytozome: a comparative platform for green plant genomics. Nucleic Acids Research 40, 1178–1186 (2011).

45. Walkowiak, S. et al. Multiple wheat genomes reveal global variation in modern breeding. Nature 588, 277–283 (2020).

46. White, B. et al. De novo annotation reveals transcriptomic complexity across the hexaploid wheat pan-genome. Nature Communications 16, 8538 (2025).

47. Ou, S. et al. Benchmarking transposable element annotation methods for creation of a streamlined, comprehensive pipeline. Genome Biology 20, 275 (2019).

48. Gurevich, A., Saveliev, V., Vyahhi, N. & Tesler, G. QUAST: quality assessment tool for genome assemblies. Bioinformatics 29, 1072–1075 (2013).

49. Simão, F.A., Waterhouse, R.M., Ioannidis, P., Kriventseva, E.V. & Zdobnov, E.M. BUSCO: assessing genome assembly and annotation completeness with single-copy orthologs. Bioinformatics 31, 3210–3212 (2015).

50. Rhie, A., Walenz, B.P., Koren, S. & Phillippy, A.M. Merqury: reference-free quality, completeness, and phasing assessment for genome assemblies. Genome Biology 21, 245 (2020).

51. Li, H. Minimap2: pairwise alignment for nucleotide sequences. Bioinformatics 34, 3094–3100 (2018).

52. Kim, D., Paggi, J.M., Park, C., Bennett, C. & Salzberg, S.L. Graph-based genome alignment and genotyping with HISAT2 and HISAT-genotype. Nature Biotechnology 37, 907–915 (2019).

53. Lovell, J.T. et al. GENESPACE tracks regions of interest and gene copy number variation across multiple genomes. eLife 11, e78526 (2022).

54. Gu, Z., Gu, L., Eils, R., Schlesner, M. & Brors, B. Circlize implements and enhances circular visualization in R. Bioinformatics 30, 2811–2812 (2014).

55. Emms, D.M. & Kelly, S. OrthoFinder: phylogenetic orthology inference for comparative genomics. Genome Biology 20, 238 (2019).

56. Baid, G. et al. DeepConsensus improves the accuracy of sequences with a gap-aware sequence transformer. Nature Biotechnology 41, 232–238 (2023).

57. Koo, D.-H. et al. Extrachromosomal circular DNA-based amplification and transmission of herbicide resistance in crop weed *Amaranthus palmeri*. Proceedings of the National Academy of Sciences of the United States of America 115, 3332–3337 (2018).

58. Koo, D.-H., Friebe, B. & Gill, B.S. Homoeologous recombination: a novel and efficient system for broadening the genetic variability in wheat. Agronomy 10, 1059 (2020).

59. do Vale Martins, L., et al. Meiotic crossovers characterized by haplotype-specific chromosome painting in maize. Nature Communications 10, 4604 (2019).

60. Adhikari, L. et al. Genetic characterization and curation of diploid A-genome wheat species. Plant Physiology 188, 2101-2114 (2022).

61. Bray, N.L., Pimentel, H., Melsted, P. & Pachter, L. Near-optimal probabilistic RNA-seq quantification. Nature Biotechnology 34, 525–527 (2016).

62. Li, H. A statistical framework for SNP calling, mutation discovery, association mapping and population genetical parameter estimation from sequencing data. Bioinformatics 27, 2987–2993 (2011).

63. Van der Auwera, G.A. & O’Connor, B.D. Genomics in the Cloud: Using Docker, GATK, and WDL in Terra, (O’Reilly Media, 2020).

64. Bradbury, P.J. et al. TASSEL: software for association mapping of complex traits in diverse samples. Bioinformatics 23, 2633–2635 (2007).

65. Paradis, E. & Schliep, K. ape 5.0: an environment for modern phylogenetics and evolutionary analyses in R. Bioinformatics 35, 526–528 (2019).

66. Chen, W.-C. PhD Thesis, Iowa State University (2011).

67. Gain, C. & François, O. LEA 3: Factor models in population genetics and ecological genomics with R. Molecular Ecology Resources 21, 2738–2748 (2021).

68. Francis, R.M. POPHELPER: an R package and web app to analyse and visualize population structure. Molecular Ecology Resources 17, 27–32 (2017).

69. Nei, M. Analysis of gene diversity in subdivided populations. Proceedings of the National Academy of Sciences of the United States of America 70, 3321–3323 (1973).

70. Danecek, P. et al. The variant call format and VCFtools. Bioinformatics 27, 2156–2158 (2011).

71. Adhikari, L. et al. Genomic characterization and gene bank curation of *Aegilops*: the wild relatives of wheat. Frontiers in Plant Science 14, 1268370 (2023).

72. Weir, B.S. & Cockerham, C.C. Estimating F-statistics for the analysis of population structure. Evolution 38, 1358–1370 (1984).

73. Purcell, S. et al. PLINK: a tool set for whole-genome association and population-based linkage analyses. Am J Hum Genet 81, 559–75 (2007).

74. Pickrell, J.K. & Pritchard, J.K. Inference of population splits and mixtures from genome-wide allele frequency data. PLOS Genetics 8, e1002967 (2012).

75. Bolger, A.M., Lohse, M. & Usadel, B. Trimmomatic: a flexible trimmer for Illumina sequence data. Bioinformatics 30, 2114–2120 (2014).

76. Kokot, M., Długosz, M. & Deorowicz, S. KMC 3: counting and manipulating k-mer statistics. Bioinformatics 33, 2759–2761 (2017).

77. Hagberg, A.A., Schult, D.A. & Swart, P.J. Exploring network structure, dynamics, and function using NetworkX. in Proceedings of the 7th Python in Science Conference (SciPy2008) (eds Varoquaux, G., Vaught, T. & Millman, J.) 11–15 (Pasadena, CA USA, 2008).

78. Klopfenstein, D.V. et al. GOATOOLS: A python library for gene ontology analyses. Scientific Reports 8, 10872 (2018).

