## Extended Data file for "Evolutionary dynamics of the Timopheevii wheat lineage"

**Extended Data Table 1.** Summary of *T. timopheevii* assembly and annotation.

|  | **TA877** | **TA2804** | **KU-1984B** |
| --- | --- | --- | --- |
| Length of HiFi assembly (bp) | 9,557,212,470 | 9,524,506,626 | 9,545,238,165 |
| Number of contigs | 3,879 | 4,649 | 4,660 |
| Contig N50 (bp) | 32,955,232 | 42,740,592 | 33,015,809 |
| Contig N90 (bp) | 4,861,799 | 5,496,836 | 5,588,789 |
| Length of hybrid assembly (bp)^1^ | 9,655,926,988 | 9,715,815,168 | - |
| Length of hybrid scaffolds (bp) | 9,445,733,231 | 9,469,235,852 | - |
| Number of hybrid scaffolds | 292 | 281 | - |
| Hybrid scaffold N50 (bp) | 142,223,090 | 153,744,984 | - |
| Length of pseudomolecule assembly (bp)^2^ | 9,656,280,388 | 9,716,225,005 | 9,545,350,565 |
| Number of anchored hybrid scaffolds and contigs | 291 | 344 | 930 |
| Number of gaps in anchored pseudomolecules | 312 | 330 | 916 |
| Total length of unanchored chromosome (bp) | 332,089,357 | 423,054,985 | 284,810,312 |
| Number of unanchored hybrid scaffolds and contigs | 3,223 | 4,080 | 3,877 |
| BUSCO scores |  |  |  |
| Complete | 99.4% | 99.5% | 99.4% |
| Single | 5.2% | 5.4% | 5.8% |
| Duplicated | 94.2% | 94.1% | 93.7% |
| Fragmented | 0.3% | 0.3% | 0.3% |
| Missing | 0.3% | 0.2% | 0.2% |
| *k*-mer based completeness | 97.9% | 98.2% | 98.5% |
| Quality value (QV) score | 64.37 | 62.57 | 66.33 |
| Number of high-confidence genes on pseudomolecules | 87,967 | 88,243 | 88,257 |

^1^Hybrid scaffolds were created by integrating optical maps into the contig-level assemblies. The hybrid assemblies include all hybrid scaffolds and contigs that were not integrated into hybrid scaffolds.

^2^Includes contig-level assembly, optical map, and Hi-C data for TA877 and TA2804, for KU-1984B it only includes contig-level assembly and Hi-C data.

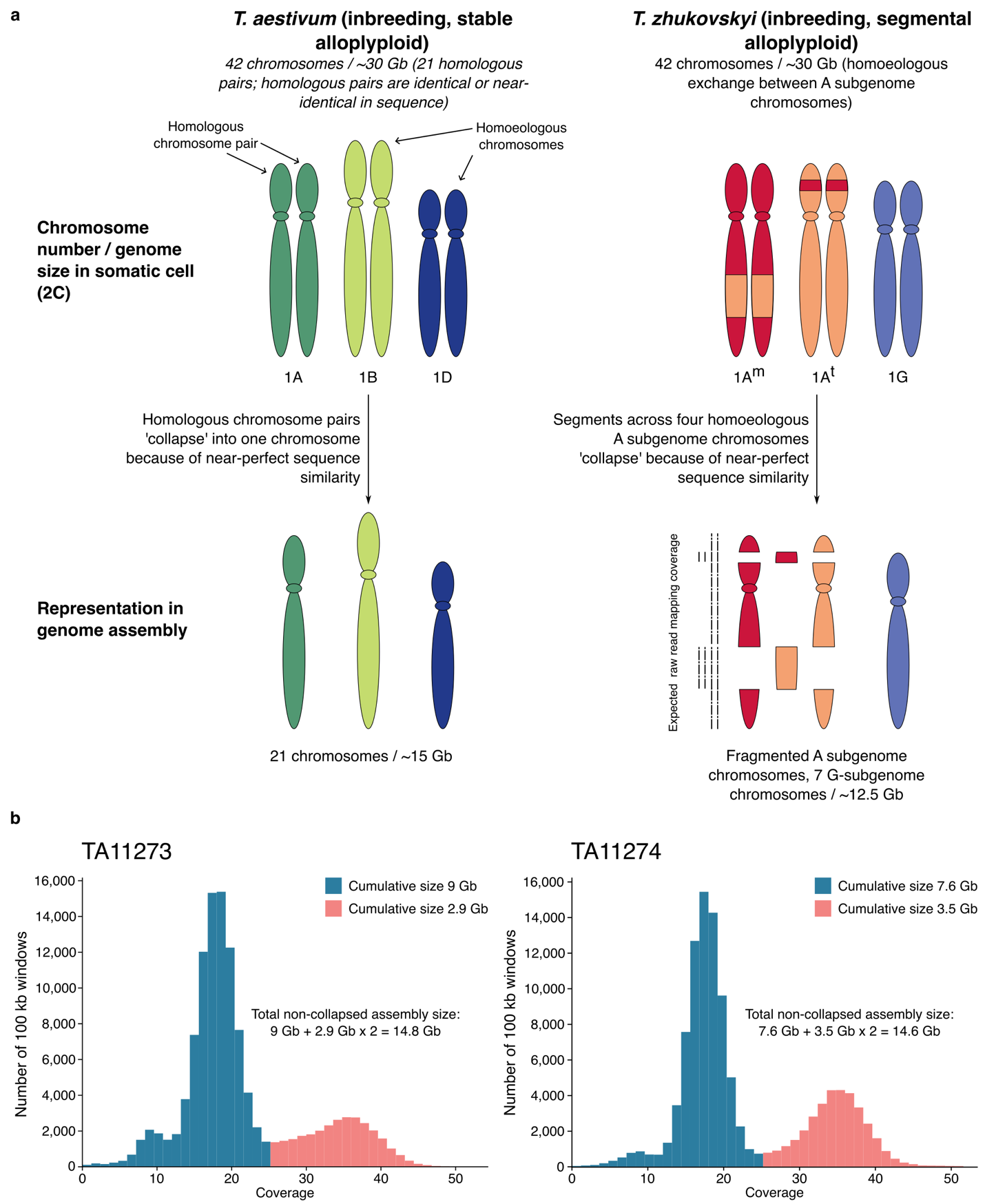

**Extended Data Fig. 1. Comparison of the assembly situations in *T. aestivum* and *T. zhukovskyi*. a**, Schematic representation of the assembly situations. Both wheat species contain 42 chromosomes (21 homologous chromosome pairs) in somatic cells (2C), with a total genome size of approximately 30 Gb. Bread wheat is inbreeding, resulting in near-identical homologous chromosome pairs. Because of this near-perfect sequence similarity, homologous chromosomes collapse during assembling, resulting in bread wheat genome assemblies generally reporting 21 chromosomes and a genome size of approximately 15 Gb. In *T. zhukovskyi*, homoeologous exchanges between the A^m^ and A^t^ chromosomes occur, resulting in chromosome segments that show near-perfect sequence similarity across all four A-genome chromosomes (i.e., having the same origin in all four chromosomes). Like the homologous chromosome pairs of bread wheat, these near-identical regions collapse in the assembly. For a segmental allopolyploid wheat species like *T. zhukovskyi*, this results in a fragmented assembly with a total assembly size of between 10.5 and 15 Gb. **b**, Coverage plots obtained by mapping raw PacBio reads to the contig-level *T. zhukovskyi* assemblies show a bimodal distribution. The second peak at ~36-fold coverage (red), approximately twice the expected depth, represents contigs that are identical across all four A-genome chromosomes and that collapsed during assembly.

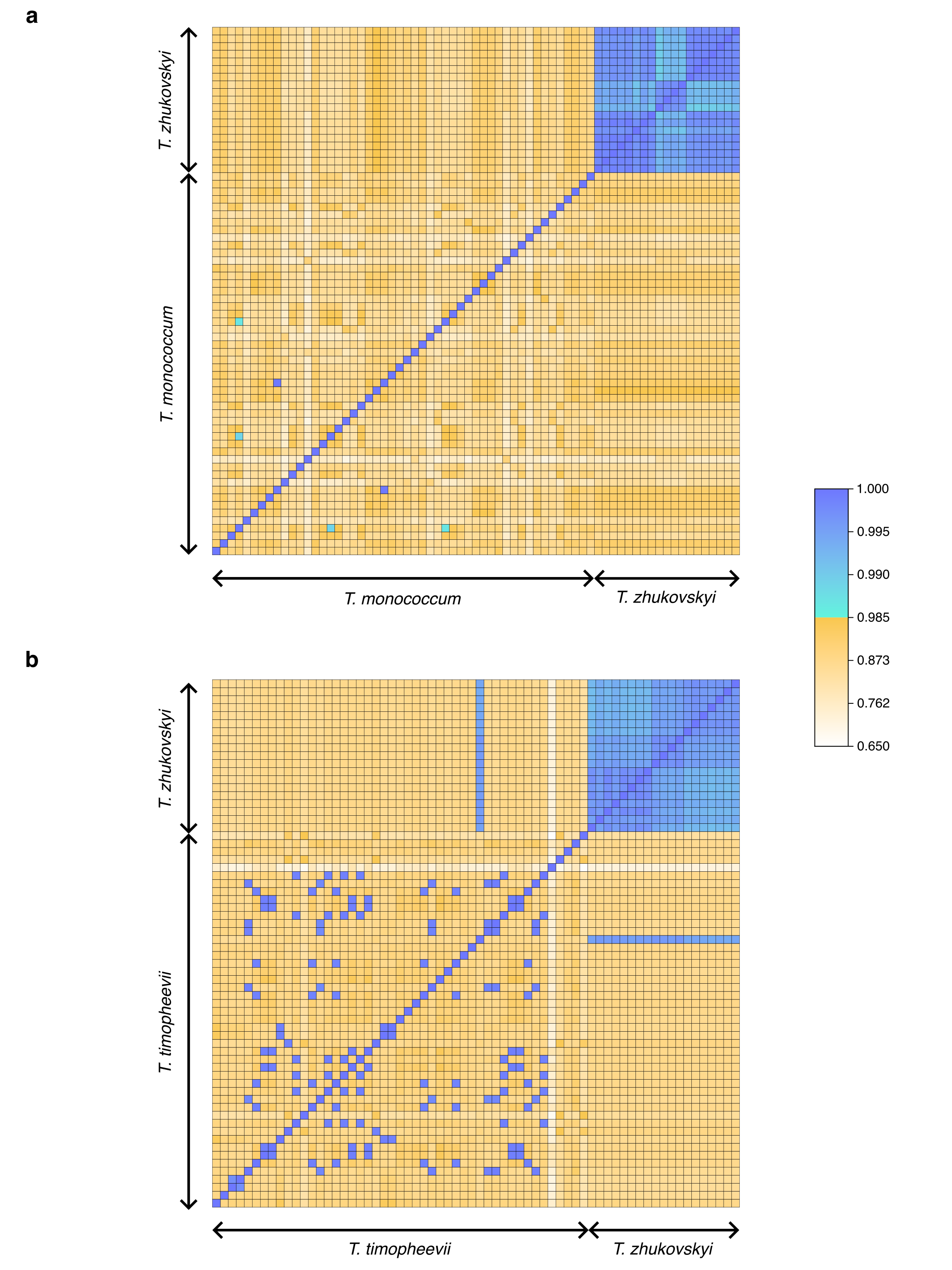

**Extended Data Fig. 2. Similarity analysis among *T. zhukovskyi* accessions. a,** Allele-matching proportional heat map for 19 *T. zhukovskyi* plants (A^m^ subgenome) and 50 domesticated einkorn (*T. monococcum* subsp. *monococcum*), based on 45,088,528 filtered sites from the seven chromosomes of *T. monococcum* (TA10622) called on the combined reference (TA10622 + TA2804). **b,** Allele-matching proportional heat map for the 19 *T*. *zhukovskyi* plants and 47 domesticated *T*. *timopheevii* ssp. *timopheevii*, using 6,142,561 filtered SNP sites from the seven chromosomes of the G subgenome called on the combined reference (TA10622 + TA2804).

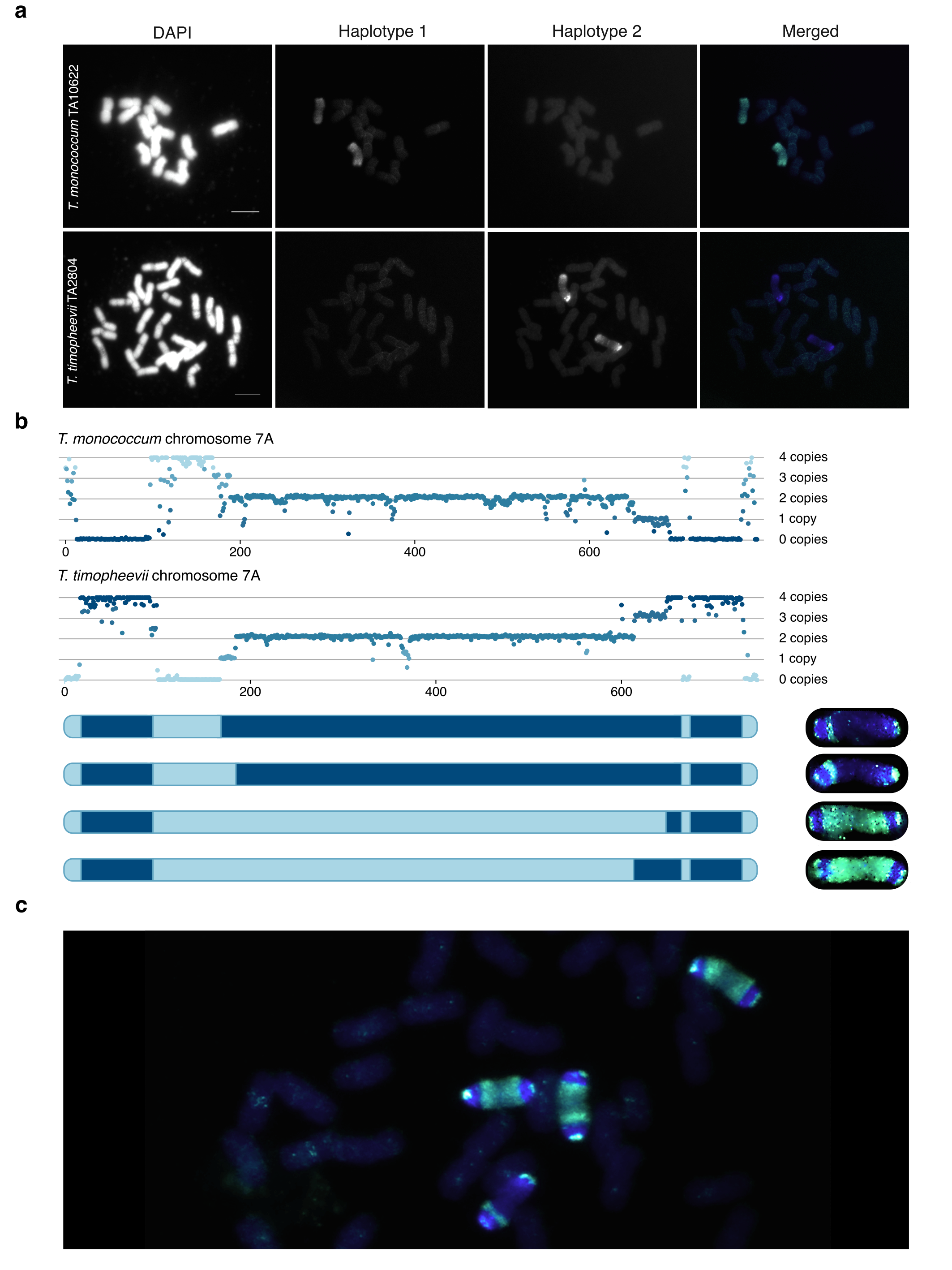

**Extended Data Fig. 3. Homoeologous exchange in hexaploid zhukovsky’s wheat. a,** Oligo painting on mitotic metaphase chromosomes of *T. monococcum* (accession TA10622) and *T. timopheevii* (accession TA2804). Probes designed on chromosome 7A of TA10622 are shown in green color and probes designed on chromosome 7A of TA2804 in blue color. **b,** Normalized read counts corresponding to chromosomes 7A^m^ and 7A^t^ in *T. zhukovskyi* plant TA11274_P3. Segments in dark blue color on the schematic chromosome representations originate from *T. timopheevii* and segments shown in light blue color from *T. monococcum* **c,** Oligo painting on mitotic metaphase chromosomes of a progeny of *T. zhukovskyi* plant TA11274_P3, indicating tetrasomic inheritance of the four A-genome chromosomes.

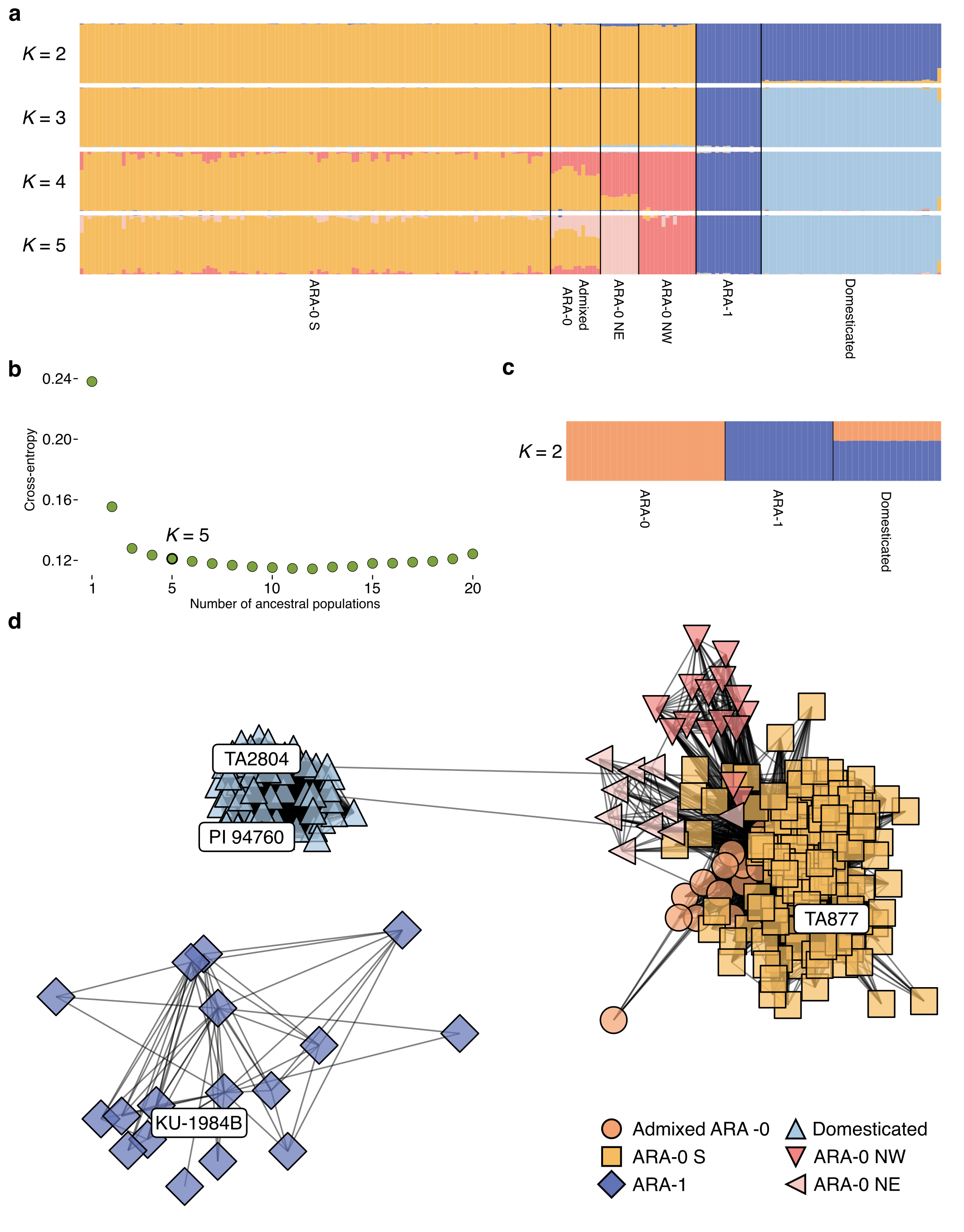

**Extended Data Fig. 4. Population structure of the *T. timopheevii* diversity panel.** **a,** Representation of the population structure from *K*=2 to *K*=5. Each vertical bar represents an accession, and the bars are filled by colors representing the proportion of each ancestry. **b,** Cross-entropy values for different values of *K*. *K*=5 is highlighted with a thicker border. **c,** Population structure representation with a balanced number of accessions from each subpopulation, revealing the admixed state of domesticated *T. timopheevii*. **d,** *k*-mer based network representation of the *T. timopheevii* population structure. Accessions with a normalized distance closer than 0.32 are connected.

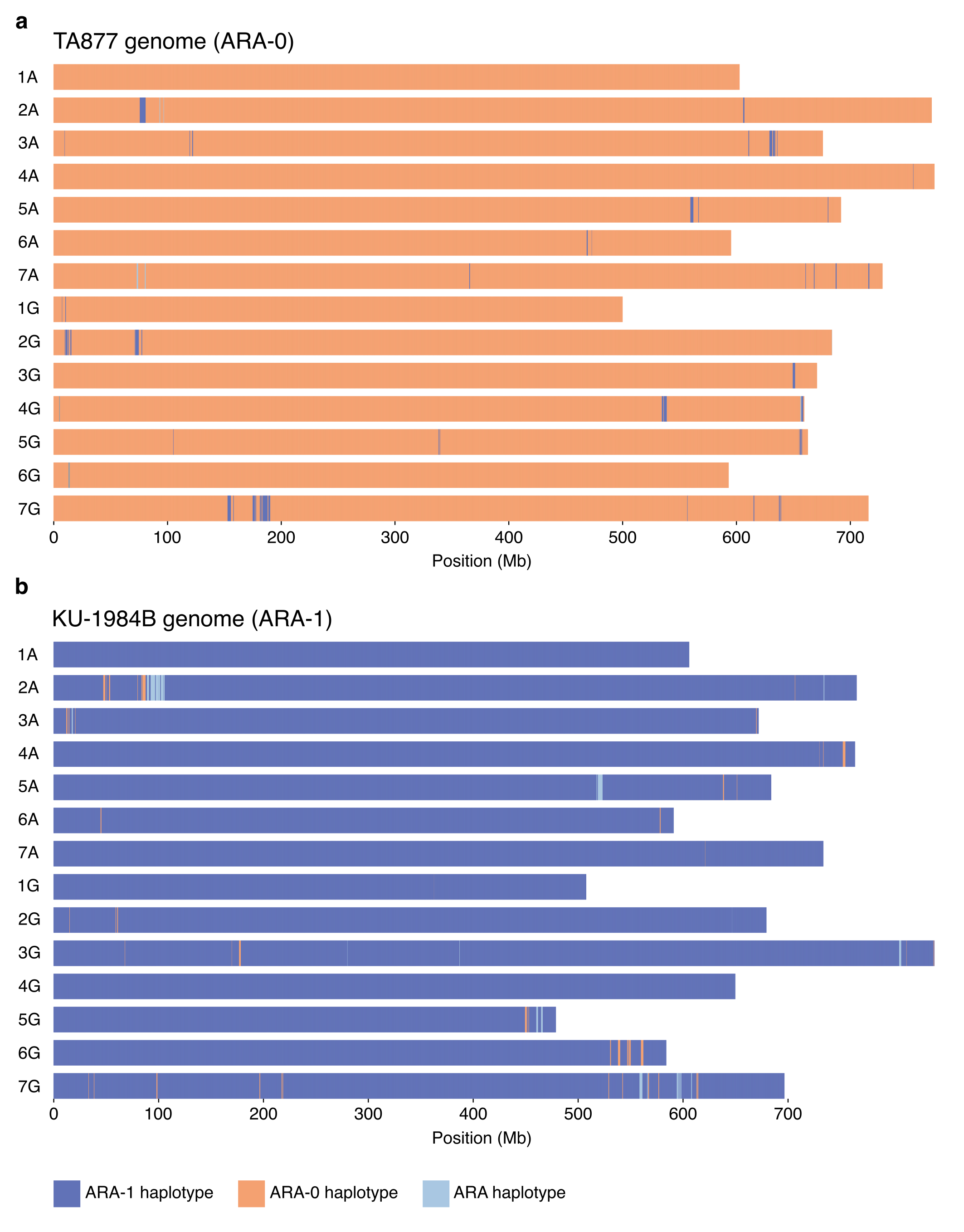

**Extended Data Fig. 5. Haplotype composition of wild *T. timopheevii.* a,** Haplotype compositions of the wild *T. timopheevii* accession TA877 considering the population structure defined for *K*=3. Shown are the chromosomes of TA877 divided into 50 kb windows. Each window is coloured according to the wild subpopulation showing identity-by-state for the respective window. **b,** Haplotype compositions of the wild *T. timopheevii* accession KU-1984B considering the population structure defined for *K*=3. Shown are the chromosomes of KU-1984B divided into 50 kb windows. Each window is coloured according to the wild subpopulation showing identity-by-state for the respective window.

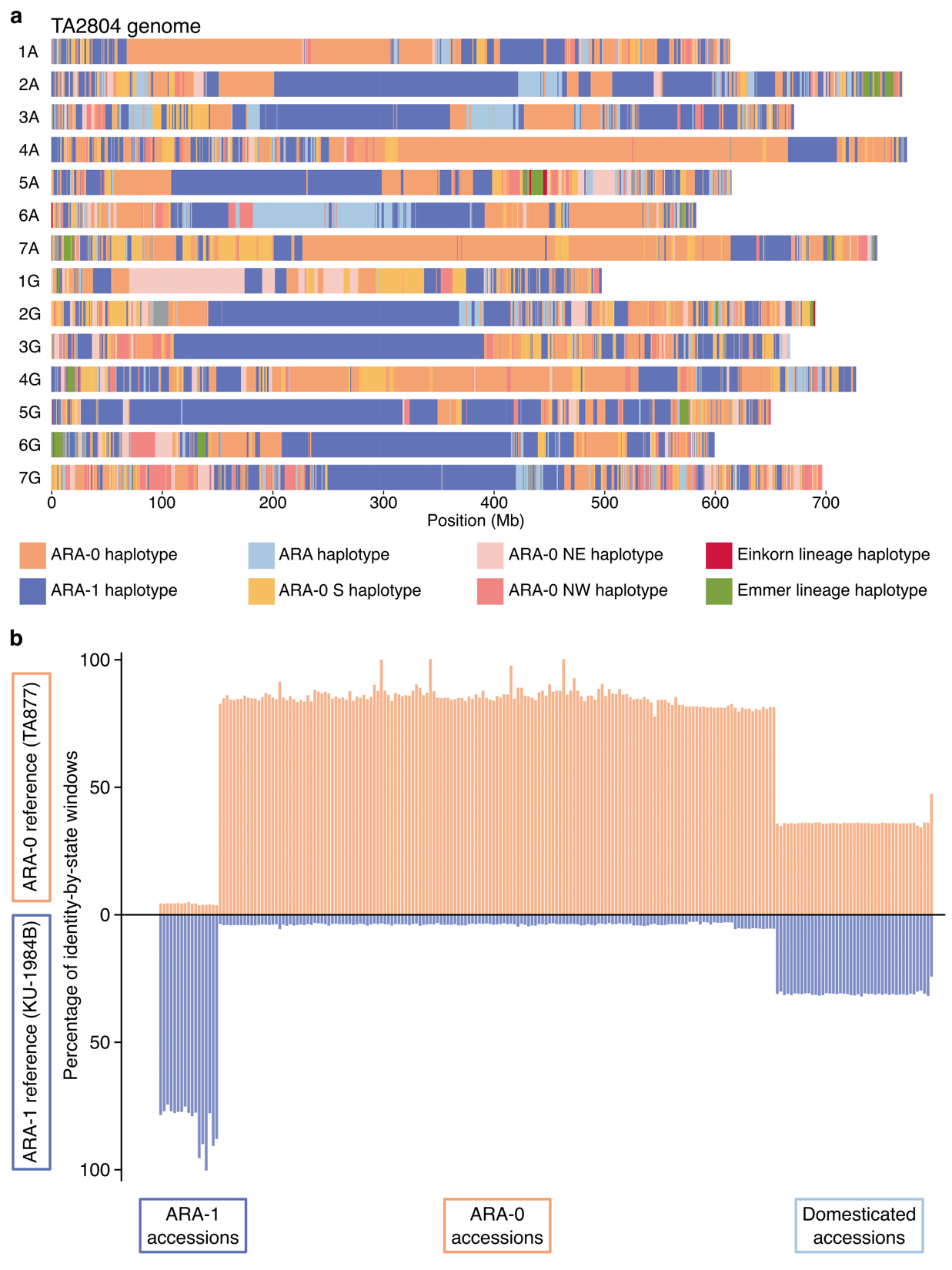

**Extended Data Fig. 6. Haplotype composition of *T. timopheevii.* a,** Haplotype compositions of the domesticated *T. timopheevii* accession TA2804 considering the population structure defined for *K*=5. Shown are the chromosomes of TA2804 divided into 50 kb windows. Each window is coloured according to the wild subpopulation showing identity-by-state for the respective window. ARA-0 indicates windows assigned to ARA-0 population, but without being able to determine the origin corresponding to one of the three ARA-0 subpopulations. **b,** Percentage of identity-by-state windows for each of the *T. timopheevii* accessions against the two wild references TA877 and KU-1984B.

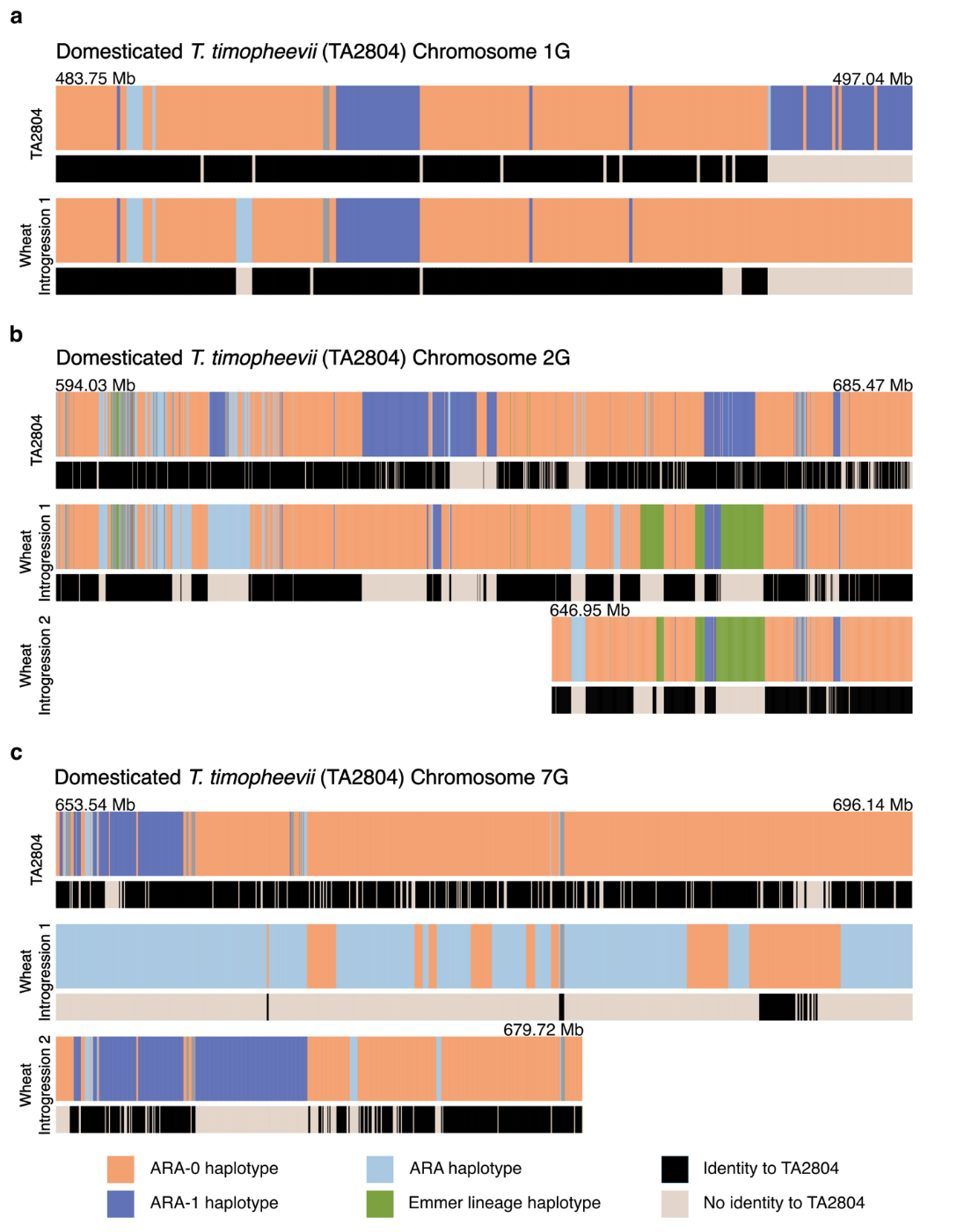

**Extended Data Fig. 7. Additional examples of *T. timopheevii* introgressions into wheat accessions of the Emmer lineage.** Shown are schematic representations of three *T. timopheevii* introgressions originating from *T. timopheevii* chromosomes 1G (**a**), 2G (**b**), and 7G (**c**). For each introgression, the topmost track corresponds to the haplotype composition of the domesticated *T. timopheevii* accession TA2804, with salmon colour indicating identity-by-state to ARA-0 and dark blue colour showing identity-by-state to ARA-1. Black bars of the second track indicates that all the domesticated *T. timopheevii* and *T. zhukovskyi* accessions included in our diversity panel showed identity-by-state across the segments, reflecting the restricted genetic diversity across extant domesticated *T. timopheevii*. The tracks labelled wheat introgressions correspond to *T. timopheevii* introgressions on chromosomes 1D (**a**), 2B (**b**), and 7B (**c**). The grey bars in the lower tracks indicate chromosome segments that are not identical-by-state to the domesticated *T. timopheevii* TA2804, most likely indicating a historic introgression from a now-extinct domesticated *T. timopheevii* population. The introgression shown in **a** is largely identical to the corresponding haplotype found in the extant domesticated *T. timopheevii* population. In contrast, the introgressions shown in **b** and **c** have a haplotype composition that is absent from the extant domesticated *T. timopheevii* population, indicating a high likelihood for originating from an extinct domesticated population.

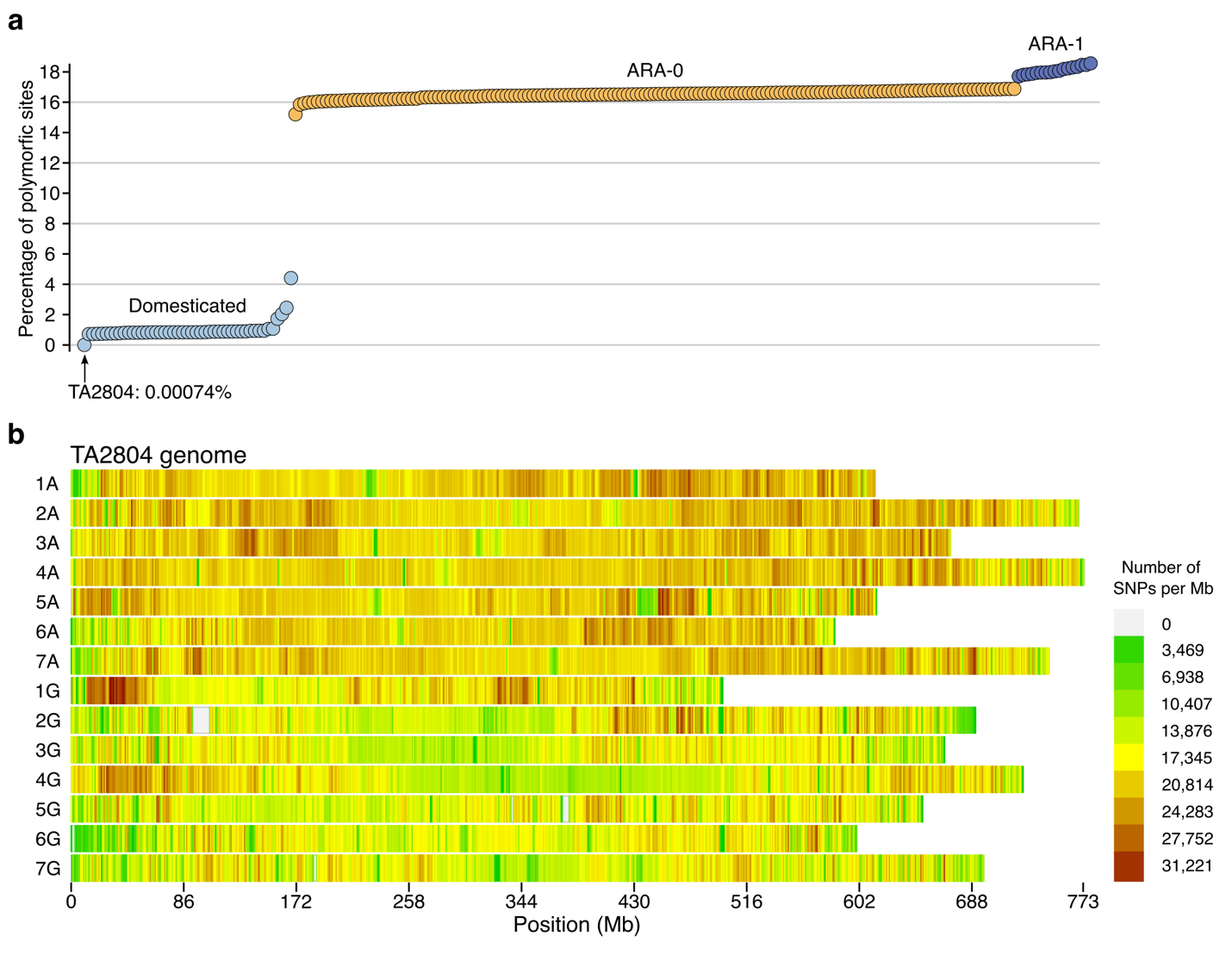

**Extended Data** **Fig. 8. SNP data statistics using TA2804 as a reference. a,** The percentage of polymorphic sites for each *T. timopheevii* accession compared to the TA2804 reference assembly. Whole-genome sequencing reads of an independently re-sequenced TA2804 plant revealed a very low error rate of 0.00074% **b,** SNP density in windows of 1 Mb computed across the 14 chromosomes of TA2804.

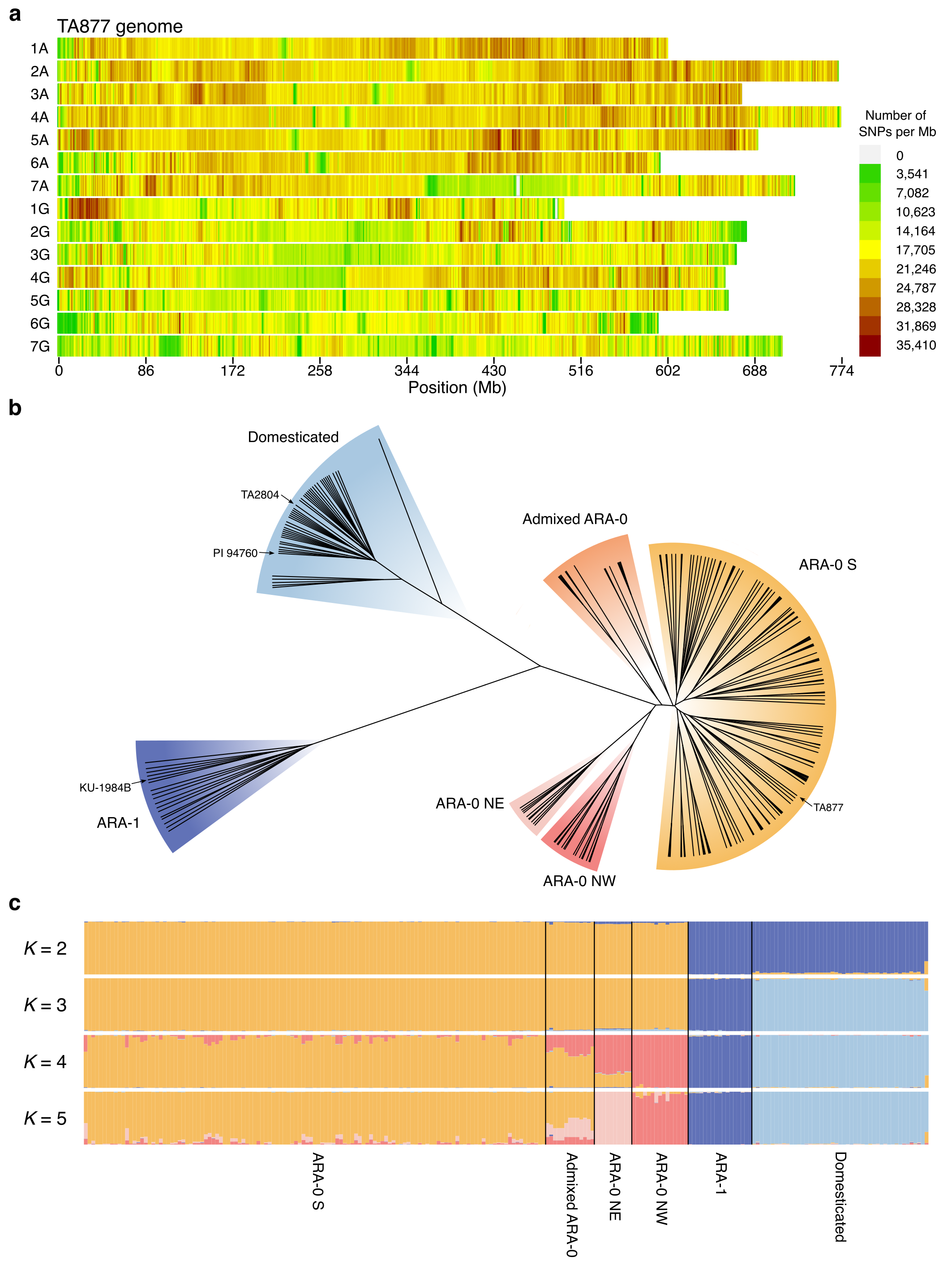

**Extended Data Fig. 9. Validation of the population structure using TA877 as reference.** Shown are the SNP density in windows of 1 Mb computed across the 14 chromosomes of TA877 (**a**), a SNP-based phylogenetic tree (**b**), and the *T. timopheevii* population structure from *K*=2 to *K*=5 (**c**).
