## Supplementary Data file for "Evolutionary dynamics of the Timopheevii wheat lineage"

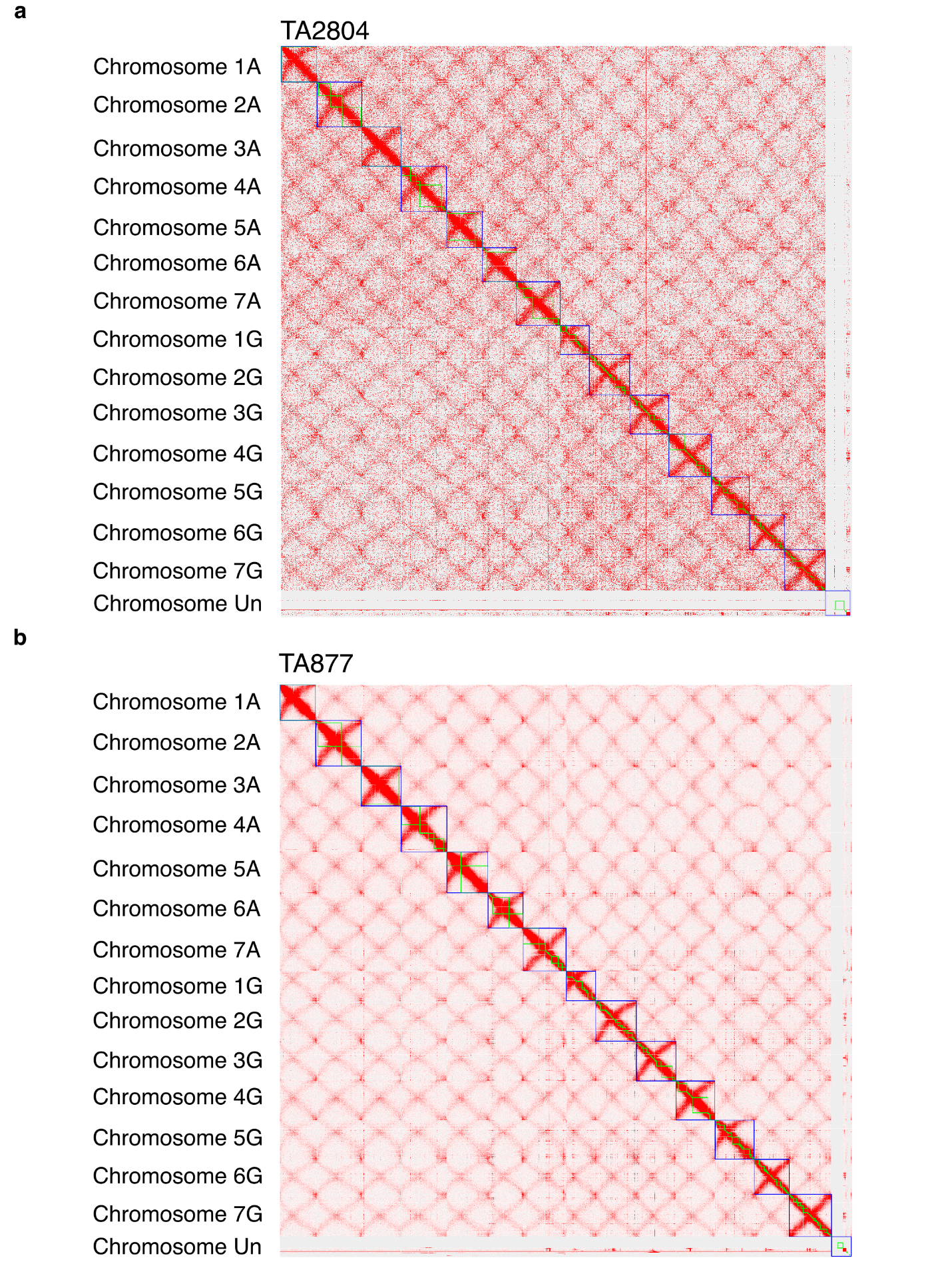

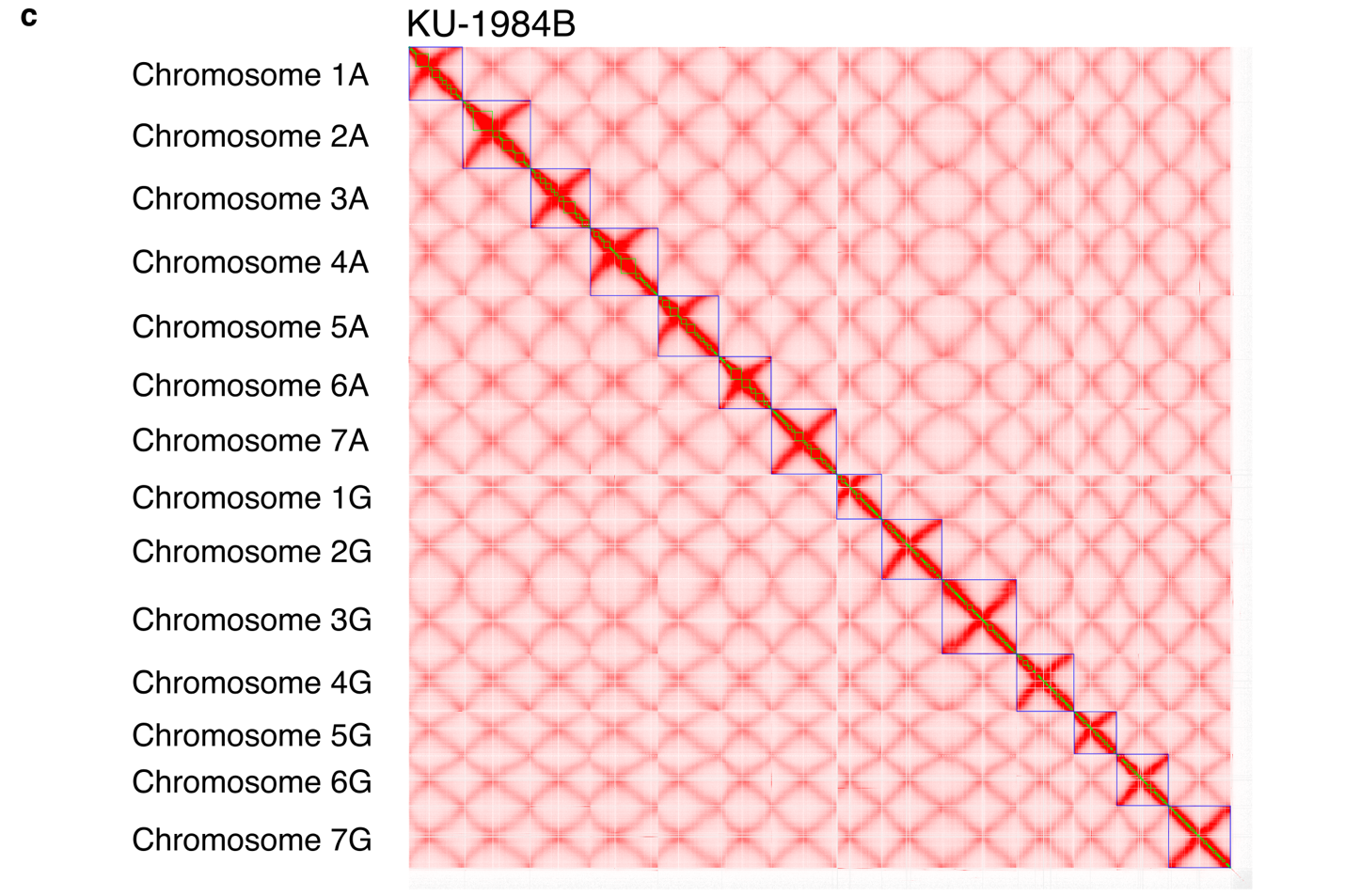

**Supplementary Fig. 1.** Chromosome contact maps of *T. timopheevii* accessions TA2804 (**a**), TA877 (**b**) and KU-1984B (**c**). Green squares indicate hybrid scaffolds, while blue squares indicate chromosomes.

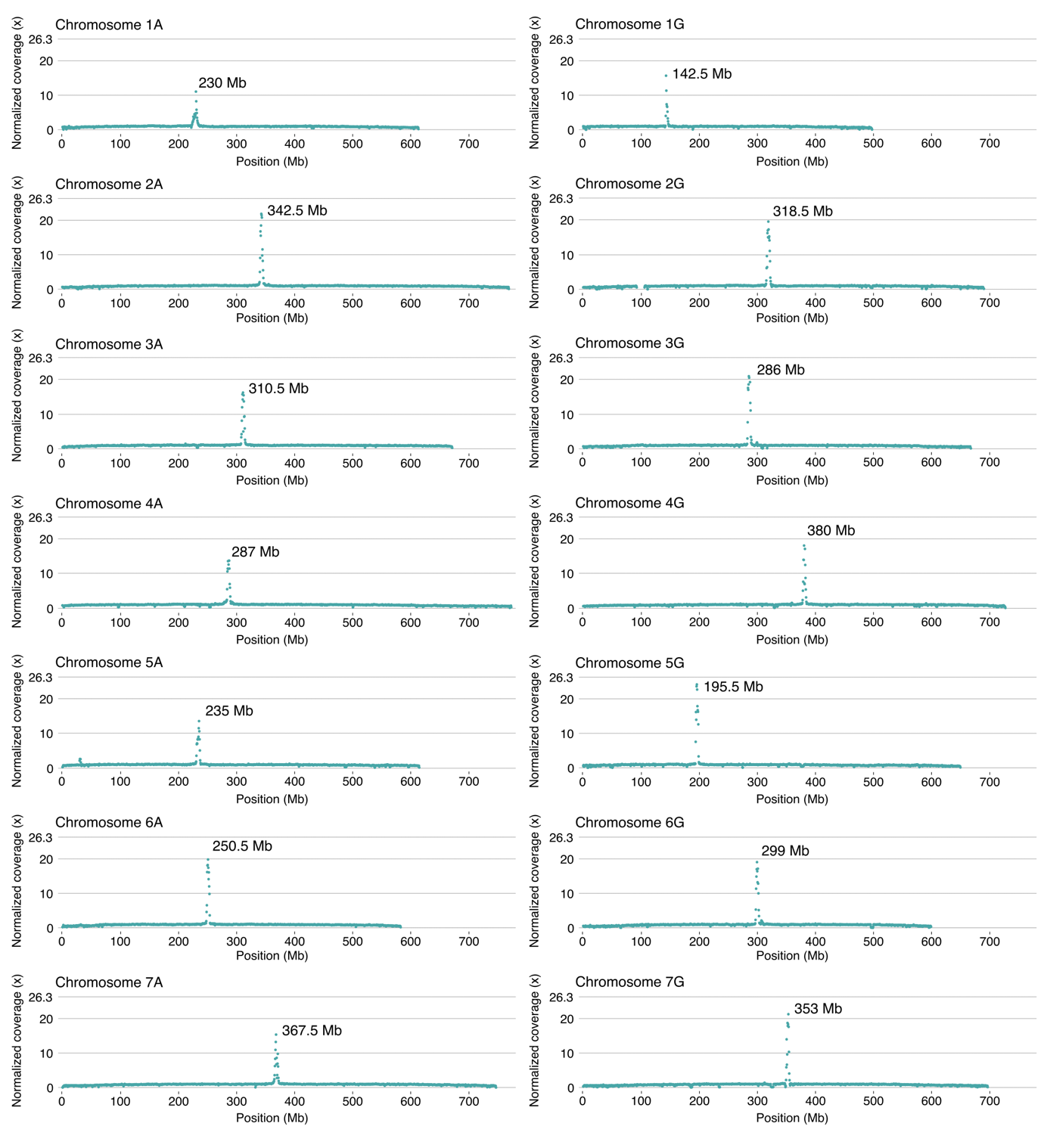
**Supplementary Fig. 2. Centromere positions in TA2804.** Shown are CENH3 ChIP-Seq read depths along the TA2804 chromosomes.

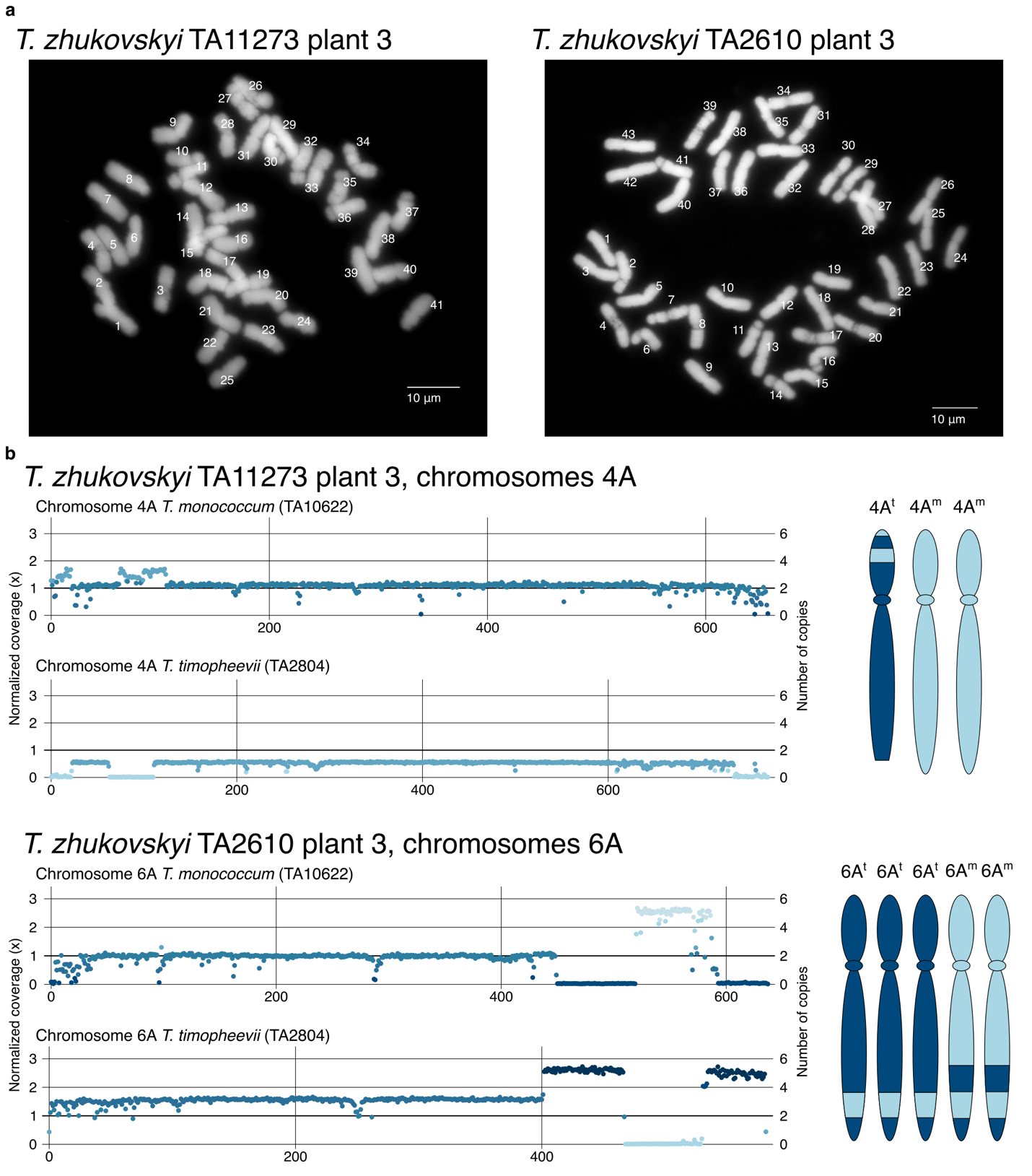

**Supplementary Fig. 3. Aneuploidy in *T. zhukovskyi.* a**, Chromosome counts in *T. zhukovskyi* plants TA11273_P3 (41 chromosomes) and TA2610_P3 (43 chromosome). **b**, Normalized read counts corresponding to chromosome 4A in TA11273_P3 and chromosome 6A in TA2610_P3. TA11273_P3 shows a total normalized read coverage of 1.5 across chromosome 4A, corresponding to three instead of four chromosome copies. Similarly, TA2610_P3 shows a total normalized read coverage of 2.5 across chromosome 6A, corresponding to five instead of four chromosome copies.

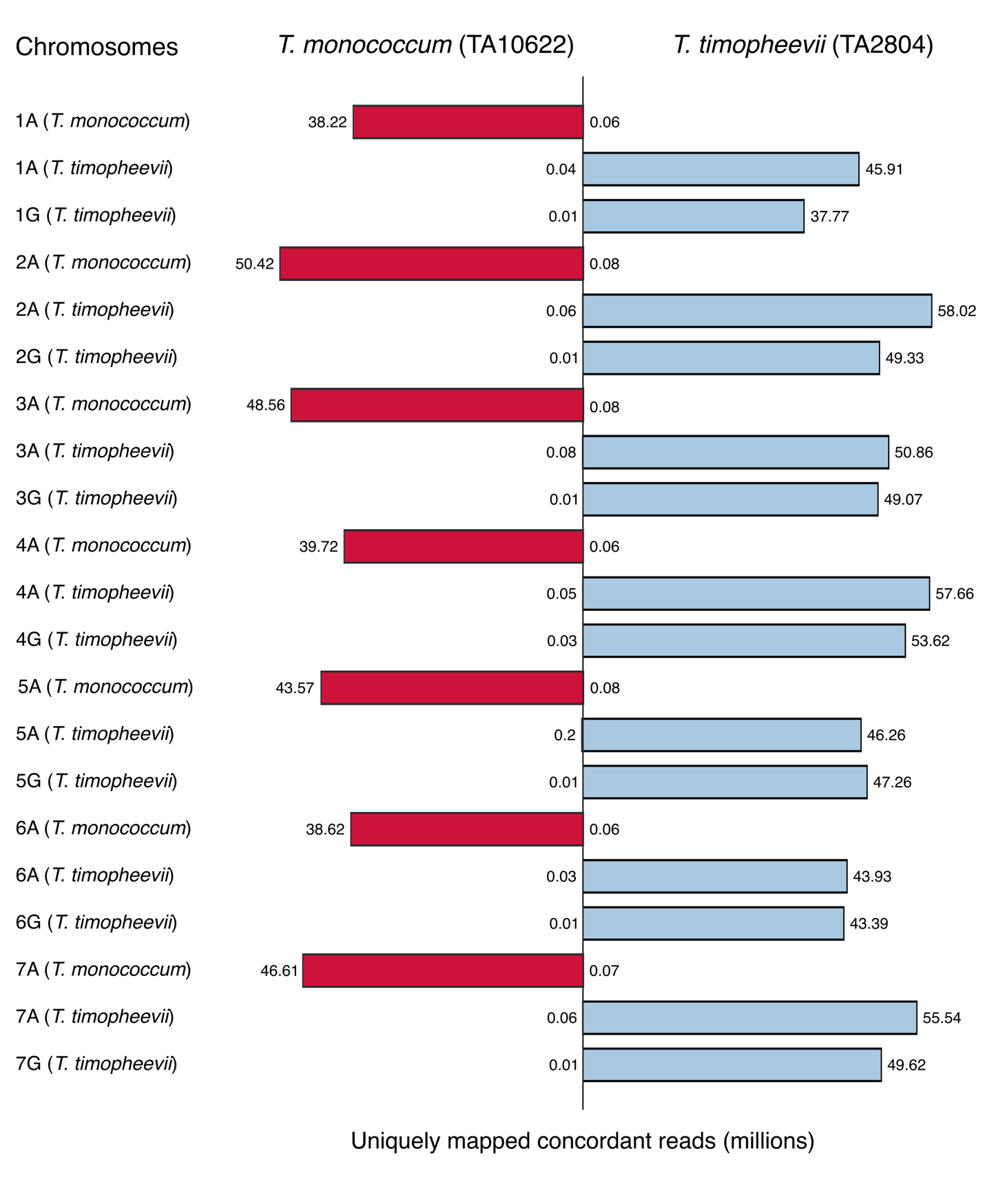
**Supplementary Fig. 4. Mapping statistics Illumina short reads from einkorn and timopheev’s wheat against a synthetic *T. zhukovskyi* (TA10622 + TA2804) reference**. The sequence reads from domesticated einkorn (TA10622) and domesticated timopheev’s wheat (TA2804) were mapped to an *in silico* synthetic *T. zhukovskyi* genome (TA10622 + TA2804). More than 99.8% of the reads mapped to their respective subgenomes. The bar chart displays the number of uniquely mapped concordant reads (in millions) across the 21 chromosomes of the *in silico* synthetic *T. zhukovskyi* genome.

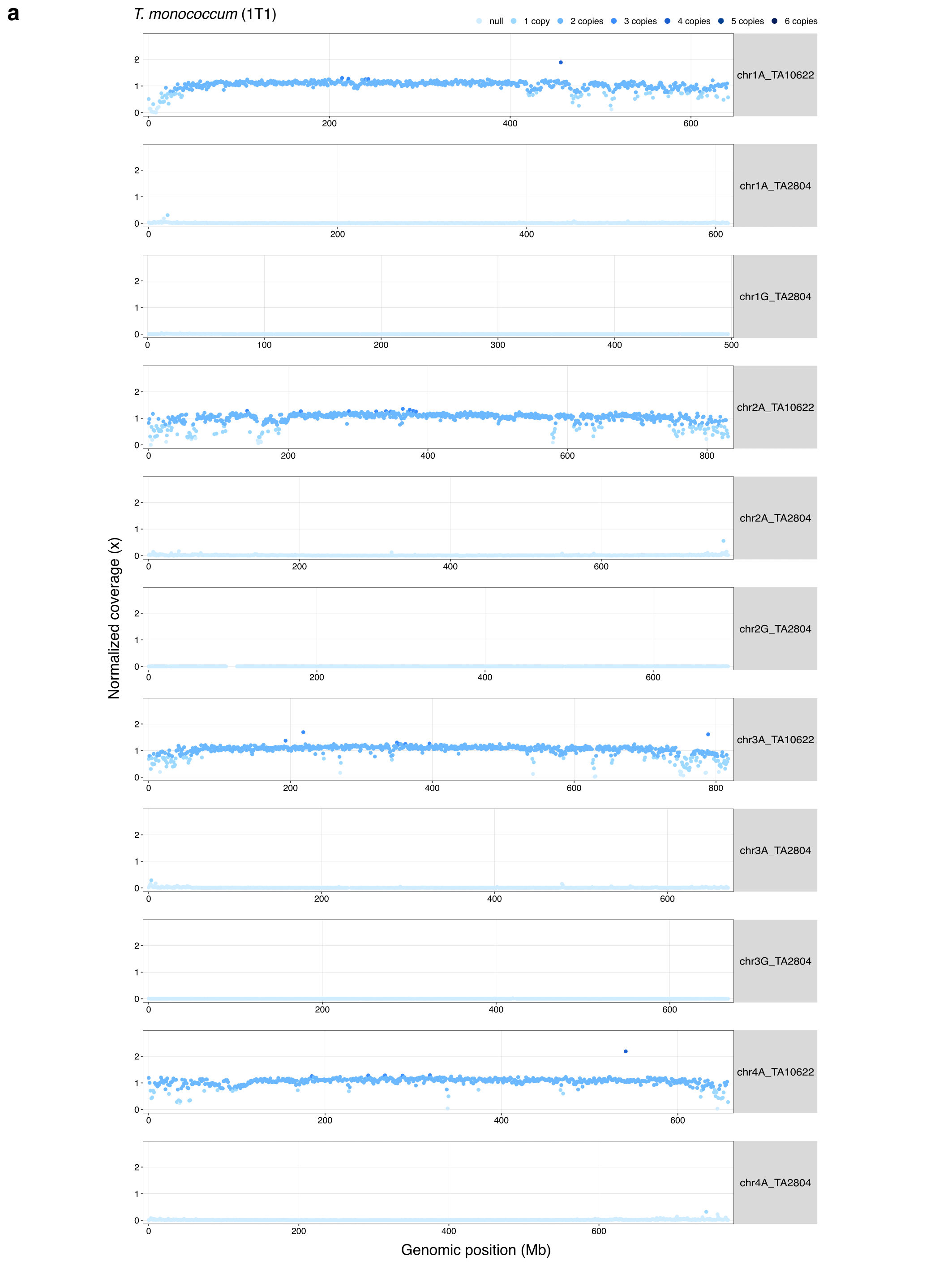

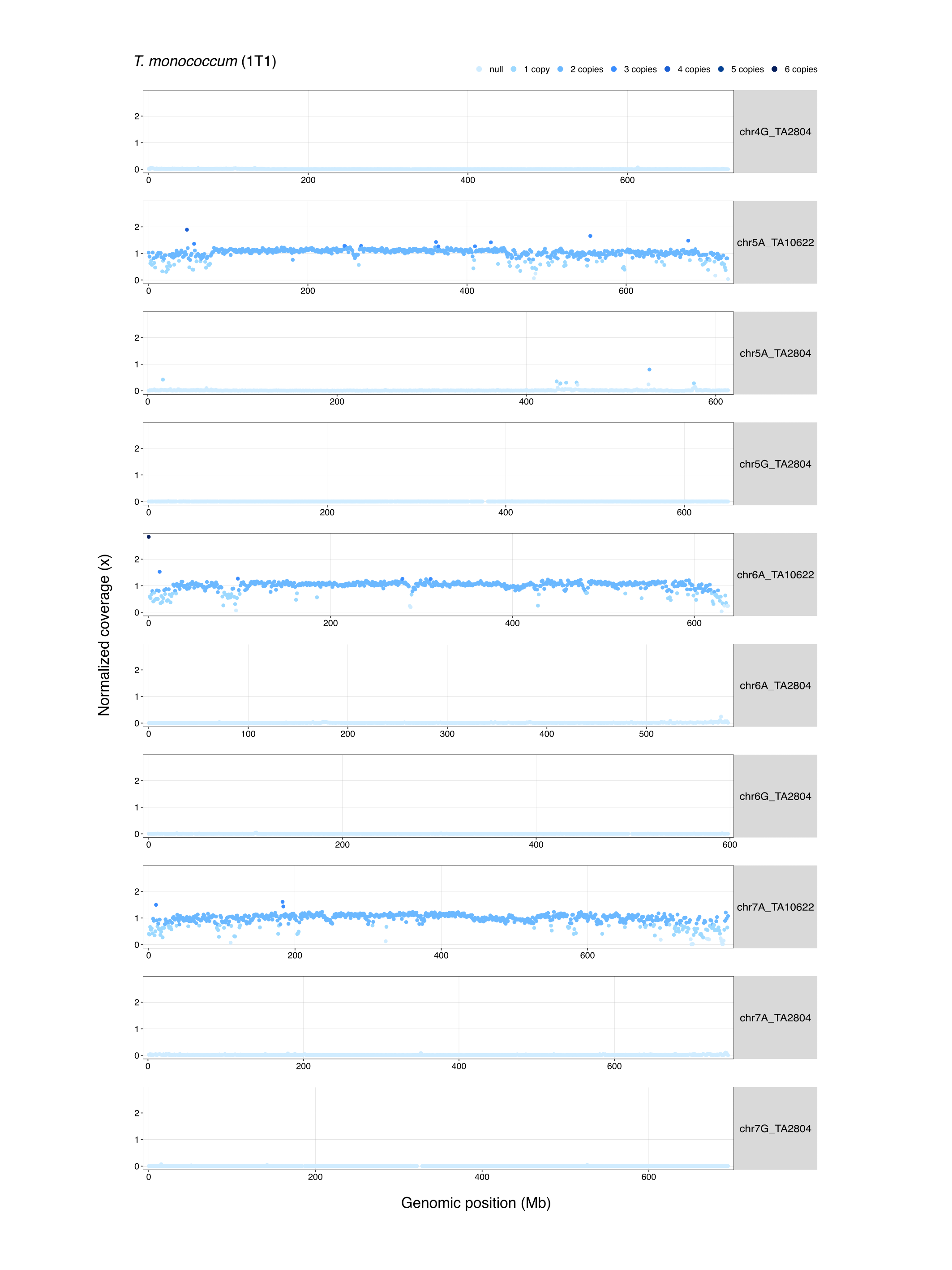

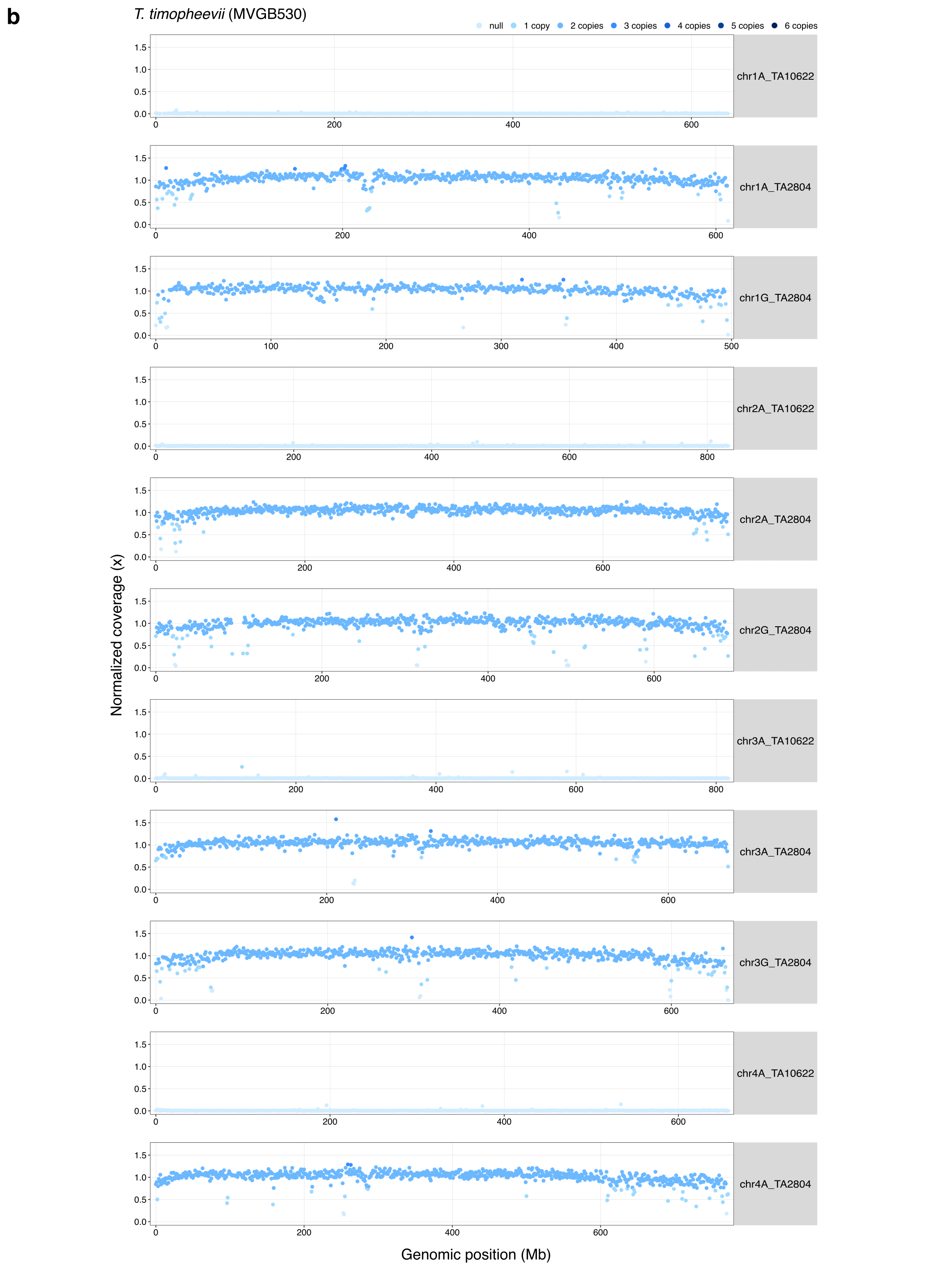

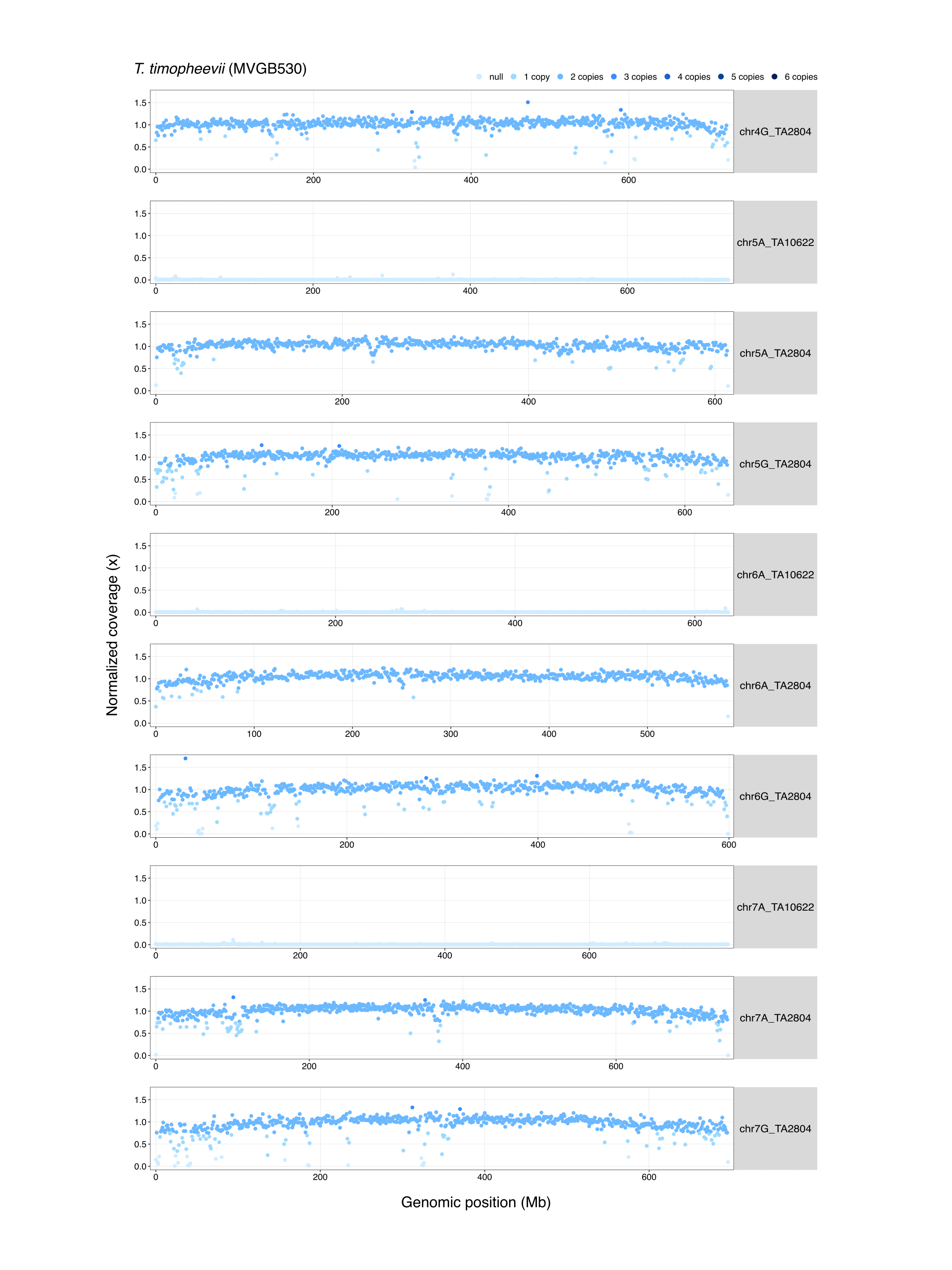

**Supplementary Fig. 5. Read counts normalized to the genome coverage indicating chromosome copy numbers.** **a,** *T*. *monococcum* accession 1T1 maps to the seven A^m^ chromosomes from TA10622. Whole-genome sequencing reads were mapped to a synthetic *T*. *zhukovskyi* (TA10622 + TA2804) reference genome, and uniquely mapped concordant reads were normalized to genome coverage **b,** *T*. *timopheevii* accession MVGB540 maps to the seven A^t^ and seven G chromosomes from TA2804.

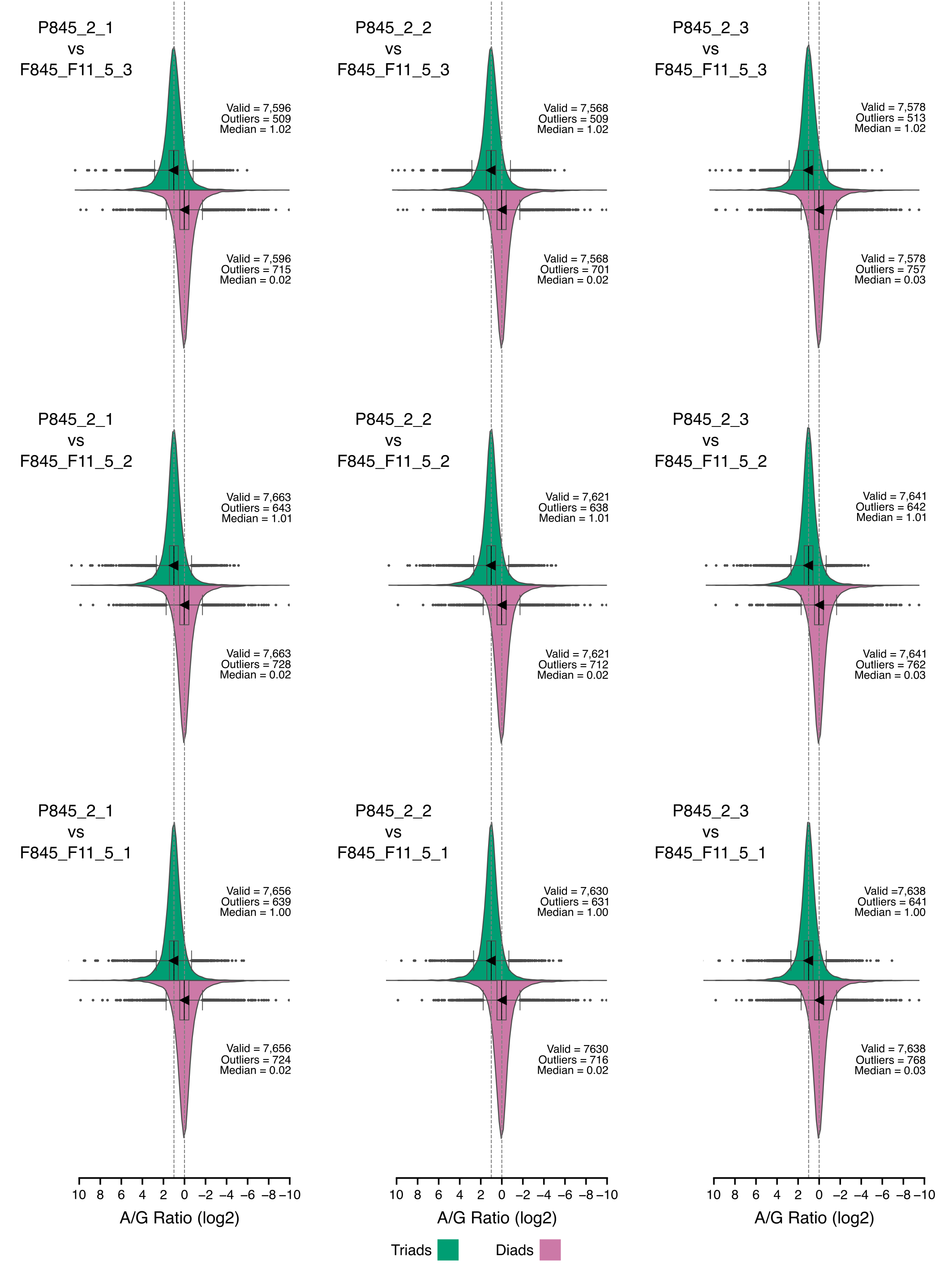

**Supplementary Fig. 6.** Violin plots showing A-to-G gene expression ratios for each pairwise comparison between tetraploid (*T. timopheevii*) in pink and hexaploid (*T. timococcum*) in green.

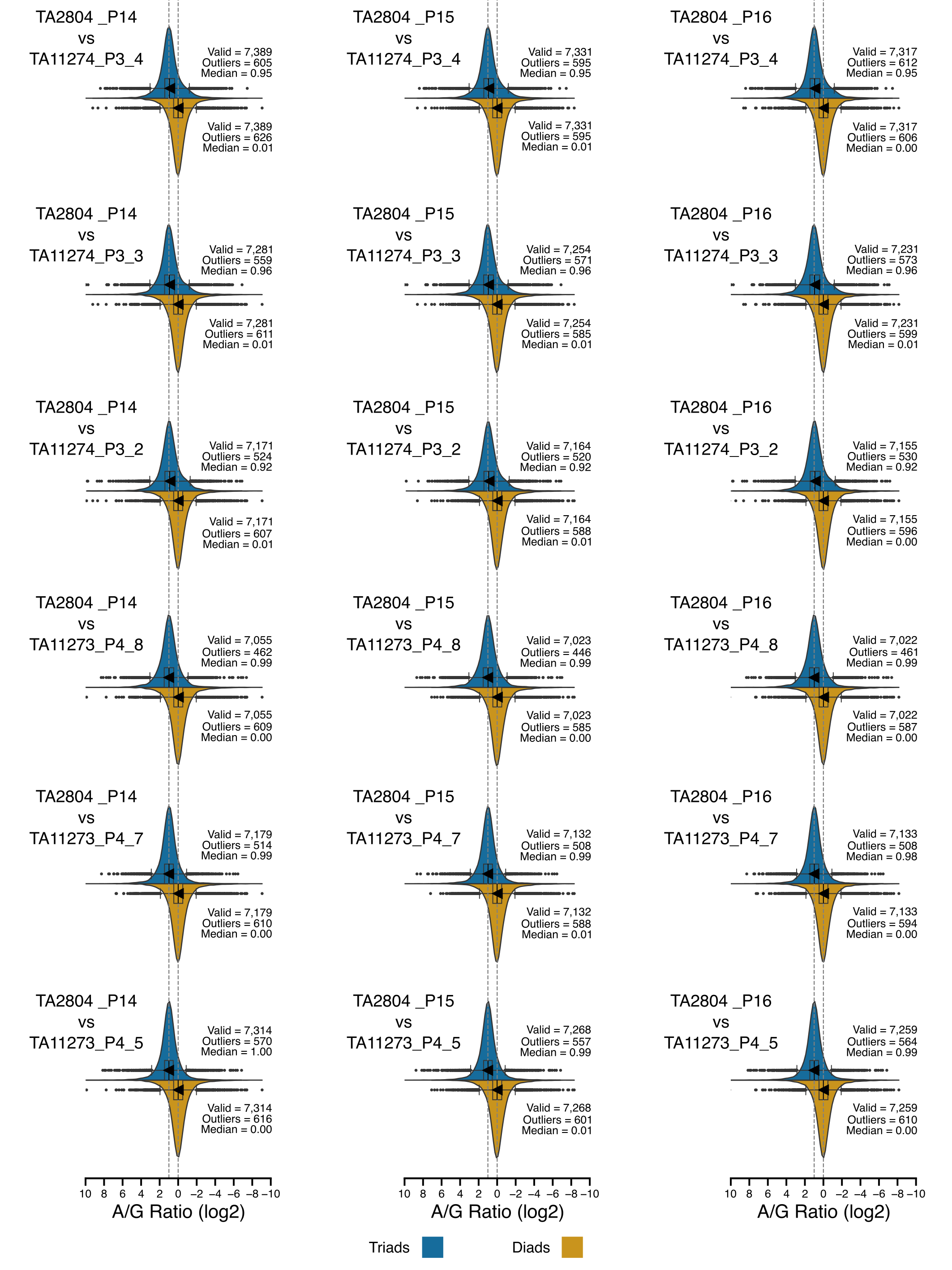

**Supplementary Fig. 7.** Violin plots showing A-to-G gene expression ratios for each pairwise comparison between tetraploid (*T. timopheevii*) in yellow and hexaploid (*T. zhukovskyi*).

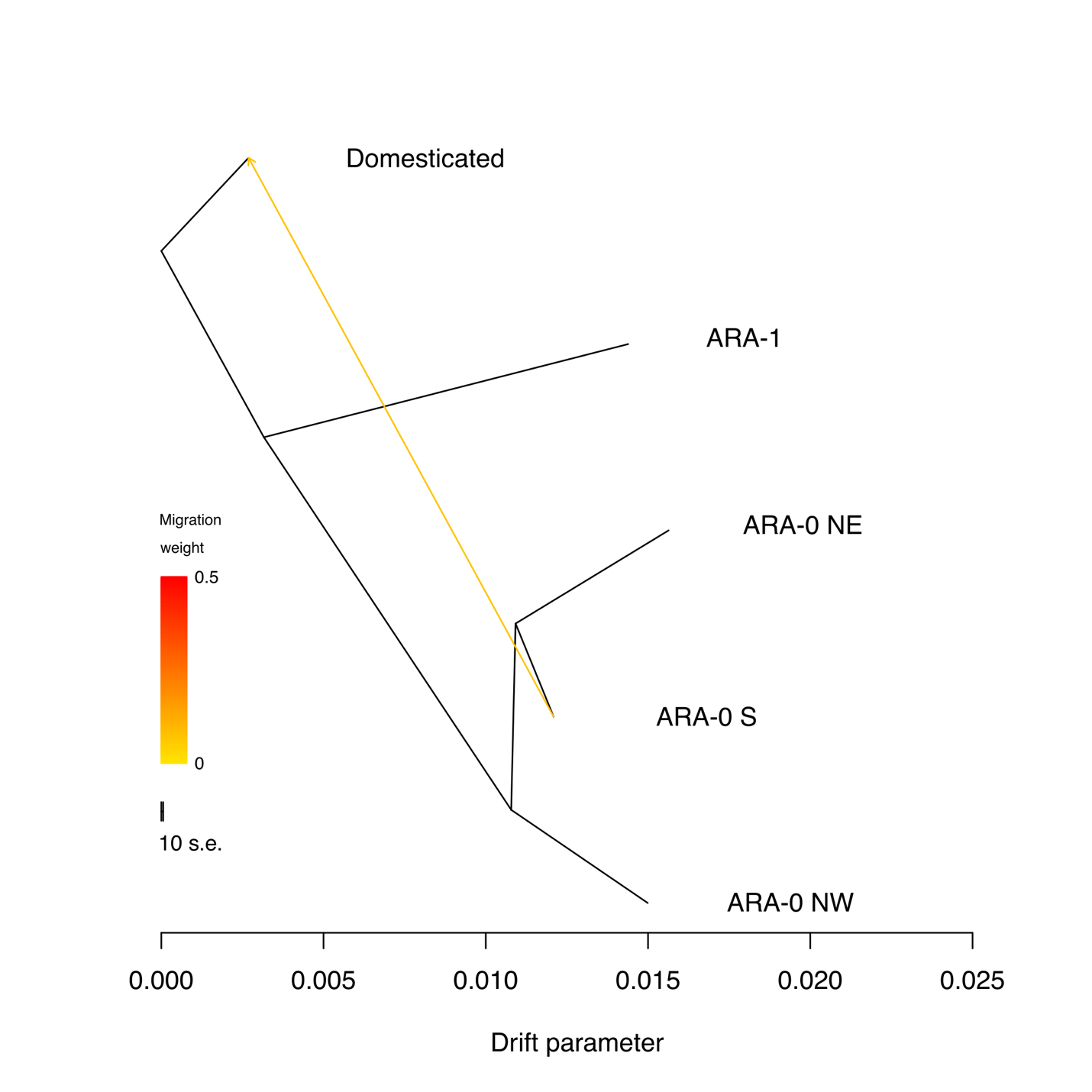

**Supplementary Fig. 8. TreeMix analysis.** TreeMix analysis ran for the five clades found in the Timopheevii wheat lineage.
