## Supplementary Notes files for "Evolutionary dynamics of the Timopheevii wheat lineage"

**Supplementary Note 1: Haplotype composition of *T. timopheevii***

This supplementary note describes how the haplotype compositions of TA2804 (Fig. 3c, Extended Data Fig. 6a), TA877 and KU-1984B (Extended Data Fig. 5a, b) were determined. The procedure includes three steps, each using a different script implemented in Python and available at https://github.com/emilecg/T.timopheevii_evolution. OpenAI's ChatGPT (https://chat.openai.com) was used to assist in refining and debugging code described here.

Step 1: IBSpy variation score distributions: The first step uses two files as inputs, (i) a single tsv file listing the variation scores obtained with FastIBS for the *T. timopheevii* accessions TA2804, TA877 and PI 94760, and (ii) a two-column tsv file listing the names of wild wheat relative accessions and their respective sub-population determined by phylogeny (Fig. 3 a, b; Extended Data Fig. 4, Supplementary Table 22).

The script computes the distribution of variation scores, with the result being a numerical description of the distribution (Figure 1 below). A separate distribution is computed for each 50-kb window of a reference assembly and for each wild wheat relative subpopulation. For example, for TA2804 (194,453 50-kb windows) and seven wild wheat relative subpopulations (four wild *T. timopheevii* subpopulations, wild emmer, *T. urartu*, wild *T. monococcum*), 1,361,171 distribution plots were generated. The bin sizes vary according to the FastIBS variation scores. Based the distribution of the variation scores across the bins, the script proceeds by defining the continuous areas present in the distribution (highlighted as a line connecting the dots in Fig. 1). For each area (peak), the start and the end of the interval, the highest peak value and the value of the area will be computed. These results are written as an output file, which will serve as the input file in the second step.

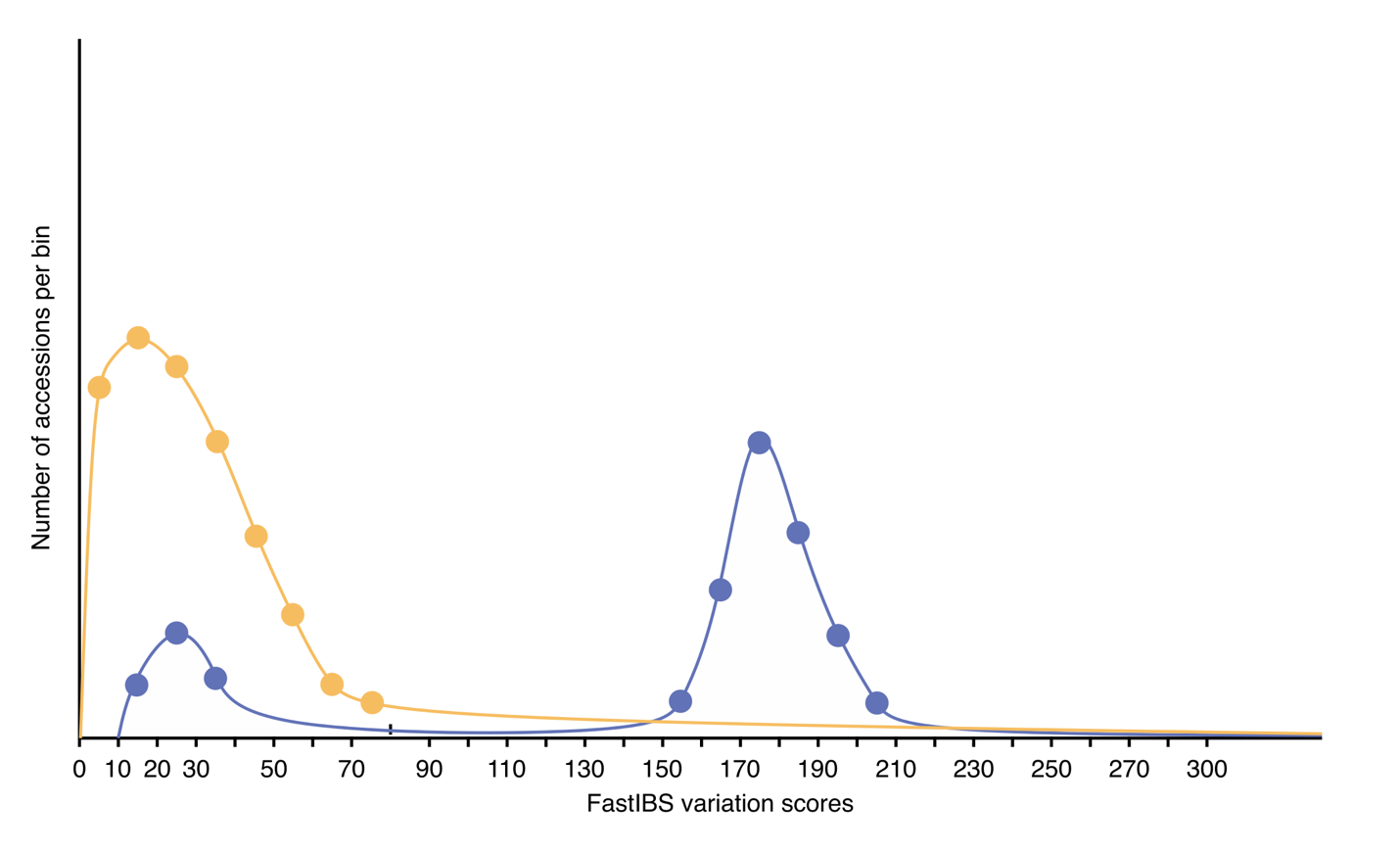

**Figure 1.** Schematic representation of the variation score distribution for two subpopulations, one in yellow and one in blue. The number of accessions per variation score bin is represented in the y-axis, while the bins (variation scores) are represented on the x-axis.

Step 2: Variation score distribution rating: In step two, a rating score will be given to each variation score distribution produced in step 1 based on the lowest variation scores and the variation score distribution (Fig. 2 below). As indicated in step 1, multiple variation score distributions are computed for each 50-kb window based on the number of wild wheat relative subpopulations.

The rating scores are assigned based on three parameters that can be tuned, (i) the minimal distance between two areas to still be considered a single area (default: 20), the variation score considered identical-by-state (default: <31), and the variation score for the interspecific range (default: 100). In total, there are eight different ratings based on the minimum variation score and the number of areas. Variation score distributions with a single peak are prioritized over multiple peak distributions because the multiple peaks might be the result of gene flow from the single peak subpopulation.

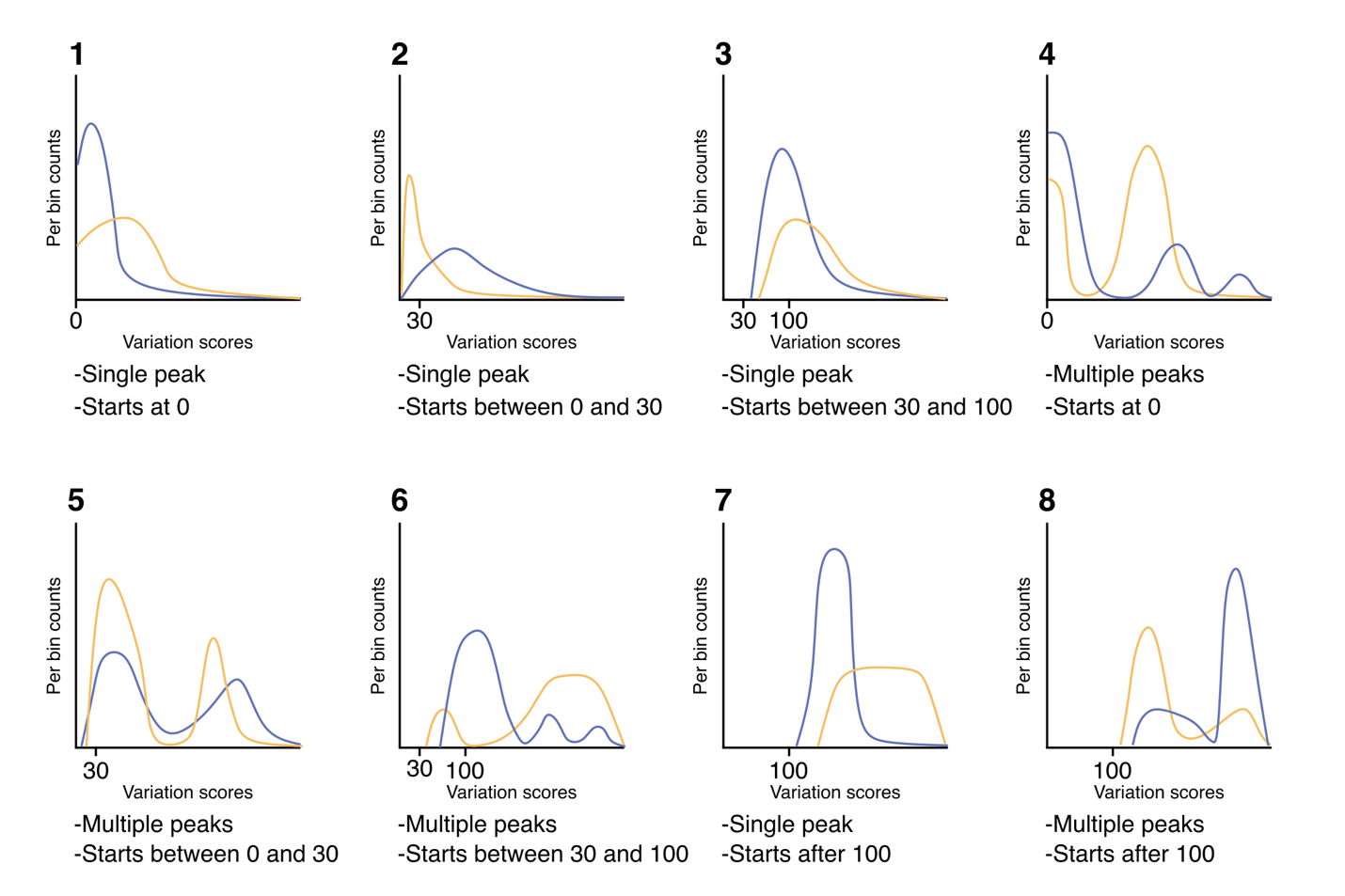

**Figure 2.** Each variation score distribution produced in step 1 is given a rating score based on the number of areas (peaks) and the minimum variation score. For each rating, two possible distribution curves are indicated in yellow and blue colour, respectively.

Step 3: Assign subpopulation origin based on rating score. For each 50-kb window of the reference assembly, the wild wheat relative subpopulation with the lowest rating score determines the subpopulation origin. If multiple subpopulations produce identical minimum rating scores with highest priority for a given 50-kb window, the selection criteria depend on the variation score distribution. For the single-area distributions, the subpopulation with the highest peak of the area being the one representing the lowest variation score is selected (Figure 3 below). For the four lowest variation sore bins, there is an error rate that is taken into consideration. These tolerance values are considered for the situations, mostly occurring in the pericentromeric region, in which two close subpopulations share the same haplotype and where the script is assigning to them in alternating pattern based on little local random variation resulting in a “hairy” kind of pattern. For multiple-areas distributions, the criterion ‘area with the minimal value representing the largest part of the accessions’ is used to select a subpopulation.

When the subpopulations having the same distribution of areas cannot be assigned even using the described criteria, the last common phylogenetic node among the subpopulations is used. For example, if it is impossible to decide between ARA-0 S and ARA-0 NE, the more general value ARA-0 will be reported.

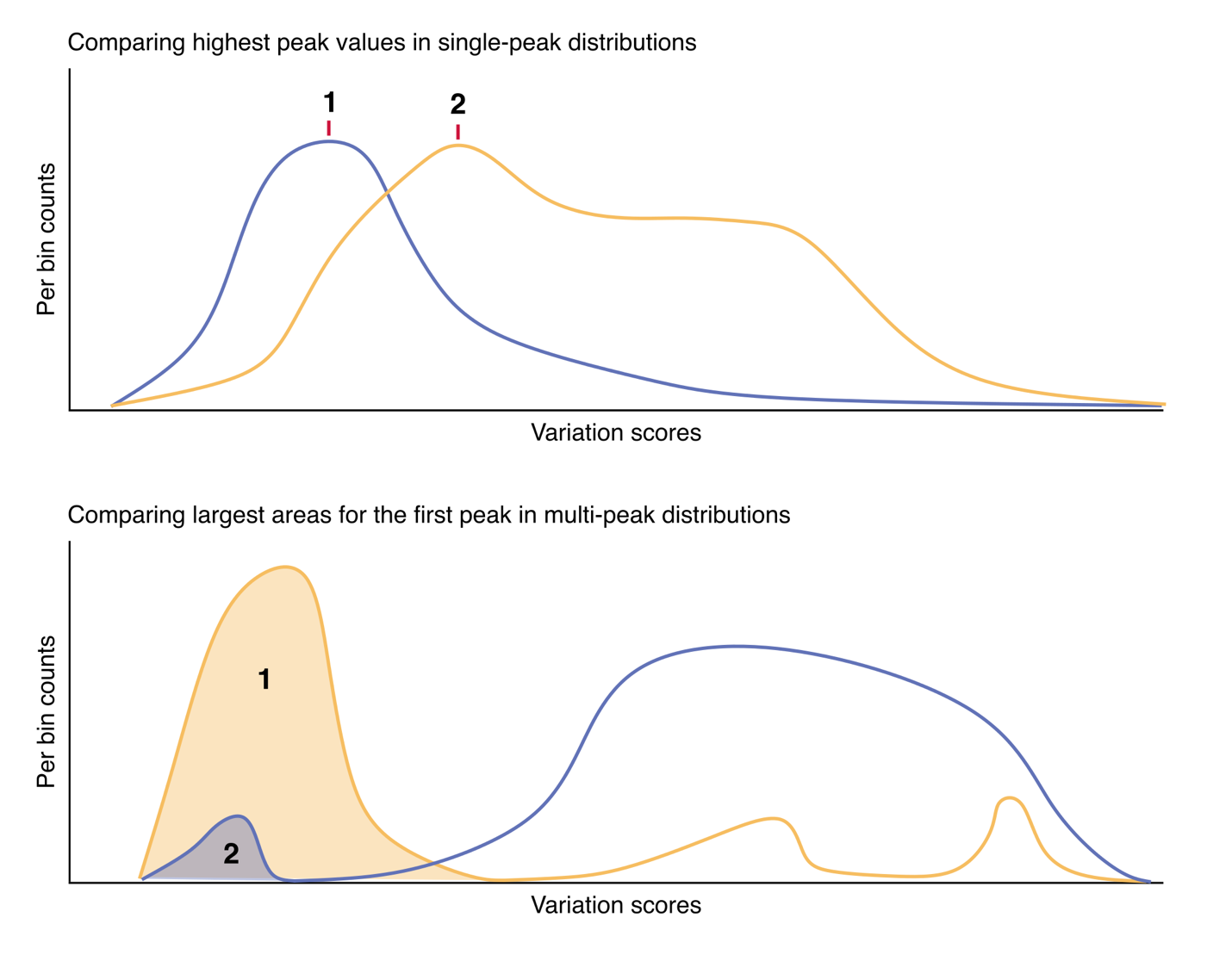

**Figure 3.** Visual representation of the selection criteria when two different subpopulations produce the same rating scores for a 50-kb window. The upper panel shows the selection criteria for a single-area distribution. The lower panel shows the selection criteria for a multiple-area distribution.

**Supplementary note 2: Introgression analysis**

1,943 accessions belonging to various domesticated Emmer wheat lineage species were used for this analysis (Supplementary Table 10). For each of the line, 31-mer sets were generated with KMC3 from trimmed FASTQ reads. The *k*-mer sets were then compared to the three reference assemblies of PI 94760, TA2804 and TA877 using FastIBS fastibs function dividing the assemblies into 50-kb windows. We decided to base the analysis mainly on TA2804, using TA877 and PI 94760 runs as a support.

The runs were converted into high-resolution heatmaps and manually examined to determine the presence of introgressions. Regions in which the TA2804 genome resulted to have introgressions from the Emmer wheat lineage were ignored.

In total, we identified 33 different *T. timopheevii* introgressions in wheat accessions of the Emmer wheat lineage. We decided to ignore introgressions occurring only in modern cultivar accessions assuming that they might derive from recent artificial introgressions. The positions and sizes of introgressions were determined based on the TA2804 genome (Supplementary Table 9) and their presence confirmed by examining the other two *T. timopheevii* references. Each introgression was named based on the chromosome arm. In cases where multiple introgressions occurred on the same chromosome arm, we used a number notation to indicate overlapping introgressions and lowercase letters to indicate non-overlapping introgressions on the same chromosome arm.

For each introgression, we identified which wheat accessions had the largest version of the respective introgression and all the wheat accessions having a partial version of it (Supplementary Table 9).

It was possible to determine the position on which the introgressions are present in the Emmer wheat lineage lines for which an assembled genome was available.

We determined high likelihood for an introgression to be a ‘new glume wheat’ type introgression by having three characteristics. The first is that wheat landraces containing the largest version of a *T. timopheevii* introgression originated from outside the modern distribution range of *T. timopheevii*. The second is that the introgression is composed by a mixture of ARA-0 and ARA-1 subpopulations, that we found to be a distinctive signature of the domesticated *T. timopheevii*. The third characteristic is that the introgression needs to be non-identical-by-state to the domesticated *T. timopheevii* sequenced in this study.

In order to establish thresholds for variation scores across species, we compared the average variation scores across different wheat species. The results of this analysis are reported in Supplementary Table 24. The thresholds obtained by this method demonstrated that the different A genomes from various wheat species are sufficiently diverged to determine introgressions based on FastIBS variation scores, particularly between the A genomes of the Timopheevii wheat lineage and Emmer wheat lineage, despite the same genome designation.
